# Combination EZH2 inhibition and retinoic acid treatment promotes differentiation and apoptosis in rhabdomyosarcoma cells

**DOI:** 10.1101/2023.06.12.544568

**Authors:** Eleanor O’Brien, Carmen Tse, Ian Tracy, Ian Reddin, Joanna Selfe, Jane Gibson, William Tapper, Reuben J Pengelly, Jinhui Gao, Ewa Aladowicz, Gemma Petts, Khin Thway, Sergey Popov, Anna Kelsey, Timothy J Underwood, Janet Shipley, Zoë S Walters

**Affiliations:** Divisions of Molecular Pathology and Cancer Therapeutics, The Institute of Cancer Research, London, United Kingdom; Cancer Sciences, Faculty of Medicine, University of Southampton, Southampton, United Kingdom; Human Development & Health, Faculty of Medicine, University of Southampton, Southampton, United Kingdom; Pathology Department, Royal Marsden NHS Foundation Trust, London, UK; Cellular Pathology Department, Cardiff and Vale UHB, Cardiff, UK; Department of Pediatric Pathology, University of Manchester Foundation Trust, Manchester, UK

**Keywords:** Rhabdomyosarcoma, Differentiation Therapy, EZH2 inhibitors, All-trans retinoic acid, Pediatric Sarcoma

## Abstract

Rhabdomyosarcomas (RMS) are predominantly pediatric sarcomas thought to originate from muscle precursor cells due to impaired myogenic differentiation. Despite intensive treatment, 5-year survival for patients with advanced disease remains low (<30%), highlighting a need for novel therapies to improve outcomes. Differentiation therapeutics are agents that induce differentiation of cancer cells from malignant to benign. The histone methyltransferase, Enhancer of Zeste Homolog 2 (EZH2) suppresses normal skeletal muscle differentiation and is highly expressed in RMS tumors. We demonstrate combining EZH2 inhibition with the differentiating agent retinoic acid (RA) is more effective at reducing cell proliferation in RMS cell lines than single agents alone. In PAX3 -FOXO1 positive RMS cells this is due to an RA-driven induction of the interferon pathway resulting in apoptosis. In fusion negative RMS, combination therapy led to an EZH2i-driven upregulation of myogenic signaling resulting in differentiation. These results provide insight into the mechanism that drives the anti-cancer effect of the EZH2/RA single agent and combination treatment and indicate that the reduction of EZH2 activity combined with the induction of RA signalling represents a potential novel therapeutic strategy to treat both subtypes of RMS.

**Highlights:**

- EZH2 expression is upregulated fusion positive (FPRMS) and fusion negative (FNRMS) rhabdomyosarcomas
- EZH2 inhibition combined with retinoic acid treatment was investigated RMS cell models.
- Combination treatment reduced cell proliferation and tumor spheroid volume.
- Combination treatment in FPRMS resulted in apoptosis in FPRMS via interferon signaling.
- Conversely, combination treatment in fusion negative RMS resulted in myogenic differentiation.

## 1. Introduction

Rhabdomyosarcoma (RMS) is a high-grade malignant tumor of mesenchymal origin that resembles skeletal muscle. RMS consists of two main subtypes: embryonal RMS (ERMS) and alveolar RMS (ARMS). ERMS comprise around 70% of cases occurring in younger children with a better prognosis, whereas ARMS accounts for up to 30% of cases, and has a poorer prognosis [1]. In ARMS, up to 80% of cases are characterized by a chromosomal translocation resulting in the formation of PAX3/FOXO1 or PAX7/FOXO1 fusion onco-proteins, key markers of poor prognoses in these cancers [2, 3]. Both ERMS and ARMS can be fusion negative (FN), and FN-ARMS have a better prognosis with outcomes similar to ERMS [4]. Although there have been incremental improvements in RMS therapy, the 5-year survival rate of patients with high-risk RMS and recurrent or metastatic disease is <30% [5], therefore there is an unmet clinical need for the identification of new therapeutic targets and strategies.

Current therapy for RMS patients involves chemotherapy, surgery, and radiotherapy; treatments that are accompanied by late side effects including reduced fertility, development issues and growth deficiency [6]. Differentiation therapy may be an alternative strategy for these patients. Activating terminal differentiation has been shown to reduce the aggressiveness of RMS by facilitating the progression to a less proliferative state [7]. Enhancer of zeste homologue 2 (EZH2) is abnormally expressed in RMS patients and cell lines [7, 8], and is associated with poor prognosis and increased metastatic potential by preventing cell differentiation whilst inducing proliferation. EZH2 is a H3K27me3 methyltransferase that forms the catalytic subunit of Polycomb repressive complex 2 (PRC2) [8, 9]. Upon activation of normal myogenesis, the levels of EZH2 decrease with the PRC2 complex dissociating from gene promoters leading to the activation of muscle-specific genes. Silencing EZH2 results in muscle differentiation through the transcriptional activation of muscle-specific promoters [10, 11]. EZH2 has also been implicated in the repression of MyoD – a core regulatory transcription factor that promotes myogenesis [11]. In fusion negative (FNRMS) tumors, EZH2 overexpression sustained proliferation [7], and EZH2 inhibition lead to myogenic differentiation [12]. EZH2 is also overexpressed in fusion positive RMS (FPRMS) tumors compared to normal muscle tissue and EZH2 depletion resulted in apoptosis in FPRMS cell models [13]. Indirect inhibition of EZH2 through PRC2 components has been shown to arrest proliferation in RMS cells, downregulating EZH2 protein levels and activity as well as global H3K27me3 levels, leading to myogenic differentiation. Together these results suggest a pro-differentiative effect of EZH2 inhibition in RMS [7, 14, 15]. However, these single agent targeting strategies did not lead to complete differentiation or apoptosis in RMS models.

Retinoic acid (RA) has been shown to inhibit proliferation and facilitate differentiation and induced apoptosis in several tumor cell lines [16–18]. The RA derivative, All-*trans* retinoic acid (ATRA) can bind and activate retinoic acid receptors (RARα, β, γ) which then regulate the expression of target genes through interactions as homodimers or as heterodimers with retinoic X receptor before binding to specific RA response elements [17]. Although RA is used in neuroblastoma to prevent recurrence as well as showing promise in the treatment of cancers such as acute promyelocytic leukemia [19, 20], RA treatment alone is not curative [21–23]. Treatment of FNRMS and FPRMS cell lines with ATRA results in a dose-dependent inhibition of cell proliferation with incomplete differentiation, suggesting that combination treatment may be required to reach terminal differentiation [24, 25] similar to acute myeloid leukemia cells where knockdown of EZH2 potentiated the pro-differentiating effects of ATRA in addition to impairing clonogenic survival [26].

Here we show that EZH2 is overexpressed in RMS patient tumors and that pharmacological inhibition of EZH2 leads to a reduction in cell proliferation and tumor spheroid volume in RMS cell models. Importantly, we demonstrate that combination treatment with EZH2 inhibitors and ATRA leads to greater efficacy than single agents alone. Finally, we link phenotypic differences seen in combination treatments in FPRMS versus FNRMS tumors with mechanistically distinct molecular changes, including the induction of interferon signaling in FPRMS tumors. This highlights the potential use of EZH2 inhibitors (EZH2i) to augment immunotherapies in these cancers for therapeutic benefit.

## 2. Materials and Methods

### 2.1. Cell culture

Cell lines, RH30, RH4, RD, RMS-01, RMS-YM, JR-1, RH41 and HFF-1 have been previously described [27]. CT-10 was a gift from Peter Houghton and HS-SY-II was a gift from Janet Shipley. CT-10 were cultured in DMEM (ThermoFisher Scientific) whereas HS-SY-II were cultured in RPMI-1640 medium (ThermoFisher Scientific) supplemented with 10% fetal bovine serum, 2 mM L-glutamine and 1% penicillin/streptomycin. Cells were maintained at 37 °C and 5% CO_2_.Cell lines were authenticated using Short Tandem Repeat fingerprinting carried out using the GenePrint 10 system (Promega). For 3D spheroids, cells were plated in ultra-low attachment (ULA) plates (ThermoFisher Scientific). The size of the spheroids was calculated by measuring two orthogonal diameters (d1 and d2). Spheroid volume was calculated using the formula: volume = 4/3πr^3^ where the radius is the geometric mean of three spheroids was calculated using r = ½√d1d2 [28].

### 2.2. Treatment of RMS cells with EZH2i or RA

Spheroids were treated every 2-3 days with either GSK343 (Sigma-Aldrich), GSK126 (Selleckchem), EPZ6438 (Selleckchem), UNC1999 (Tocris Bioscience), or UNC2400 (Tocris Bioscience) alone or in combination with ATRA (Sigma-Aldrich) as indicated. For 2D combination studies in 384-well plates the Echo 550 liquid handler (Labcyte) was programmed to add drug.

### 2.3. Immunohistochemistry (IHC)

At indicated time-points, spheroids were collected and washed in PBS before fixing in 4% paraformaldehyde for 24h at 4°C. Approximately 10 spheroids were embedded in 1% agarose, before processing as previously published [27]. Primary antibodies used were: Cleaved Caspase-3 (9661; Cell Signaling Technology (CST)), H3K27me3 (ab6002; Abcam), Ki67 (M7240; Agilent Dako), MYOG (M3559; Agilent Dako), MHC (MAB4470; R&D Systems). Spheroid cores were scored by pathologist Prof Anna Kelsey.

#### 2.3.1 EZH2 expression in tissue microarray cores

EZH2 expression was analyzed in a large cohort of RMS patient samples (n=282) and correlated with clinicopathological features. Tissue microarray slides were processed as above and incubated with EZH2 antibody (Leica Biosystems) for 1hr by Dr Frances Daley (Pathology Core Facility, ICR). Each core was scored as previously described [27]by two histopathologists blinded to patient outcomes (Dr Sergey Popov and Dr Khin Thway). Ethical approval was obtained from the Local Research Ethics Committee protocol 1836 and UK Multi-Regional Research Ethics Committee 98/4/023 (16/11/06).

### 2.5. Cell Viability and Proliferation Assay

CyQUANT cell proliferation assay (ThermoFisher Scientific) was used to measure cell proliferation in 96-well plates according to the manufacturer’s instructions. Fluorescence was measured after 1h incubation at 37°C using excitation at 485nm and emission at 530nm. To measure cell viability in 384-well plates alamarBlue cell viability reagent (ThermoFisher Scientific) was used following manufacturer’s instructions. After incubation for 4h at 37°C absorbance was measured at 570nm. Cell viability for 3D spheroids was assayed using CellTiter-Glo® 3D Cell Viability Assay (Promega) according to manufacturer’s instructions. The luminescence of each well was measured after 30min.

### 2.6. Caspase Activity Assay

Apoptosis was measured by evaluating the activation of caspase 3/7 using the Ac-DEVD-AMC Caspase-3 Fluorogenic Substrate (BD Biosciences) according to manufacturers’ instructions. The fluorescence of each well was measured after 2h incubation at 37°C at excitation at 380nm and emission at 460nm.

### 2.7. Western Blot

Cell Lysis Buffer (CST) was used to extract protein lysates and the Subcellular Protein Fractionation Kit for Cultured Cells (ThermoFisher Scientific) was used for fractionation according to manufacturers’ instructions. The following primary antibodies were used: EZH2 (5246; CST), GAPDH (MAB374; Merck Millipore), Histone H3 (ab1791; Abcam), H3K27me3 (ab6002; Abcam), MYOG (556358; BD Biosciences), p16 (ab108349; Abcam), p21 (ab80633; Abcam), PARP (9542; CST), RARα (E6Z6K; CST), α-Skeletal Muscle Actin (ab28052; Abcam), α-Tubulin (SC-8035; Santa Cruz Technology). Immunostained bands were detected by chemiluminescence (GE Healthcare).

### 2.8. RNA-sequencing (RNA-seq)

RNA was extracted from RMS cells treated with GSK343, ATRA or combination after 6 days (n=6 replicates). RNA was sent to Novogene (Hong Kong) for library preparation and sequencing. RNA-seq fastq files for each condition were aligned to the GRCh38.p13 GENCODE human genome reference using STAR (v2.7.10b) [29] with the –quantMode option set to GeneCounts. The resulting gene counts were used to identify differentially expressed genes between each condition and DMSO using Deseq2 [30]. A gene was determined as differentially expressed if the fold change between DMSO and condition was greater than two and the adjusted p-value (Benjamini-Hochberg) was less than 0.05.

### 2.9. Chromatin immunoprecipitation (ChIP) / ChIP-sequencing (ChIP-seq)

RMS cells were prepared for ChIP using SimpleChIP™ kit (#9006A; CST) according to manufacturer’s instructions. Pulldown was performed using magnetic beads and all antibodies were purchased from CST: anti-IgG (2729), anti-Histone H3 (4620), anti-H3K27me3 (9733), anti-EZH2 (5246), anti-RARα (E6Z6K). Library preparation and ChIP-Seqencing was performed by Novogene (Hong Kong). ChIP-seq fastq files for EZH2, RARα, H3K27me3, and IgG were aligned to the GRCh38.p13 human genome reference using Bowtie2 [31] and resulted BAM files sorted using Samtools (v 1.16.1) [32]. Peak calling was performed with the BAM files using MACS (v 3.0.0b1). Regular peak calling options were used for RARα, while broad peak calling options were used for EZH2 and H3K27me3, using IgG as a control. For RARα narrow peak calling the option -q 0.01 was used, and for EZH2/H3K27me3 broad peak calling the option --broad-cutoff 0.1 was used to set significance threshold. R packages ChIPseeker and TxDb.Hsapiens.UCSC.hg38.knownGene were used to annotate the called peaks. Genes for which peaks were called in or close to were compared pairwise between experiments to identify overlaps.

### 2.10. Pathway analysis and GSEA

Pathway analysis was performed on differentially expressed genes (DEGs) alone and for DEGs that peaks were called at in the ChIP-seq analysis. R packages enrichR and clusterProfiler were used to identify enriched pathways using the MSigDB hallmark genesets. Using R packages clusterProfiler and fgsea, GSEA was performed on all genes in the Deseq2 output, ranked by fold change (highest to lowest), considering MSigDB hallmark, and gene ontology (GO) genesets.

### 2.11. Statistical Analysis

Graphs represent means ± standard deviation (SD) from multiple independent experiments as stated in figure legends. Statistical significance was measured by Two-tailed unpaired t-test or One Way ANOVA as specified and assigned as follows: * *p*<0.05, ** *p<*0.01, *** *p*<0.001, **** *p*<0.0001 using GraphPad Prism. To assess synergy between treatment combinations, the Bliss predicted response (𝑌𝑎𝑏) was first calculated according to the following equation: 𝑌𝑎𝑏 = 𝑌𝑎 + 𝑌𝑏 − 𝑌𝑎𝑌𝑏 where 𝑌𝑎 and 𝑌𝑏 are observed responses with the two compounds alone. Excess over Bliss (EOB) score is calculated by subtracting the predicted response (𝑌𝑎𝑏) from the observed response of the combination treatment (𝑦𝑎𝑏) as follows: EOB = 𝑦𝑎𝑏 − 𝑌𝑎𝑏. EOB scores that are >0 represent synergy and <0 represents antagonism [33]. One-sided Fisher’s exact test was used to determine whether the overlap of genes near called peaks was significant between two different ChIP-Seq experiments. The total number of genes considered in calculations was all pseudo- and coding-genes used for annotation (n = 34,130).

## 3. Results

### 3.1. EZH2 is overexpressed in RMS primary tumors and cell lines

As EZH2 has been shown to be overexpressed in small cohorts of RMS patients [7, 13, 34], we sought to examine expression in a large, well-curated dataset of RMS patients (n=282) by tissue microarray (TMA). EZH2 was expressed in 79.4% (224/282) of patients, (Table 1; Figure 1A). Contrary to previous reports in smaller sample sets, we found no correlation between EZH2 protein expression and outcome or metastases in our cohort. The frequency and intensity of EZH2 staining was higher in the RMS sections than normal tissue samples. Variability in EZH2 staining in tumors was observed so EZH2 transcript levels was analyzed from two gene expression datasets of RMS patients [35, 36], and a dataset of childhood cancer cell lines which was compared to a normal skeletal muscle dataset using R2 [37]. *EZH2* transcript levels were higher in RMS patient samples and cell lines compared to normal skeletal muscle (Figure 1B).

**Figure 1.**
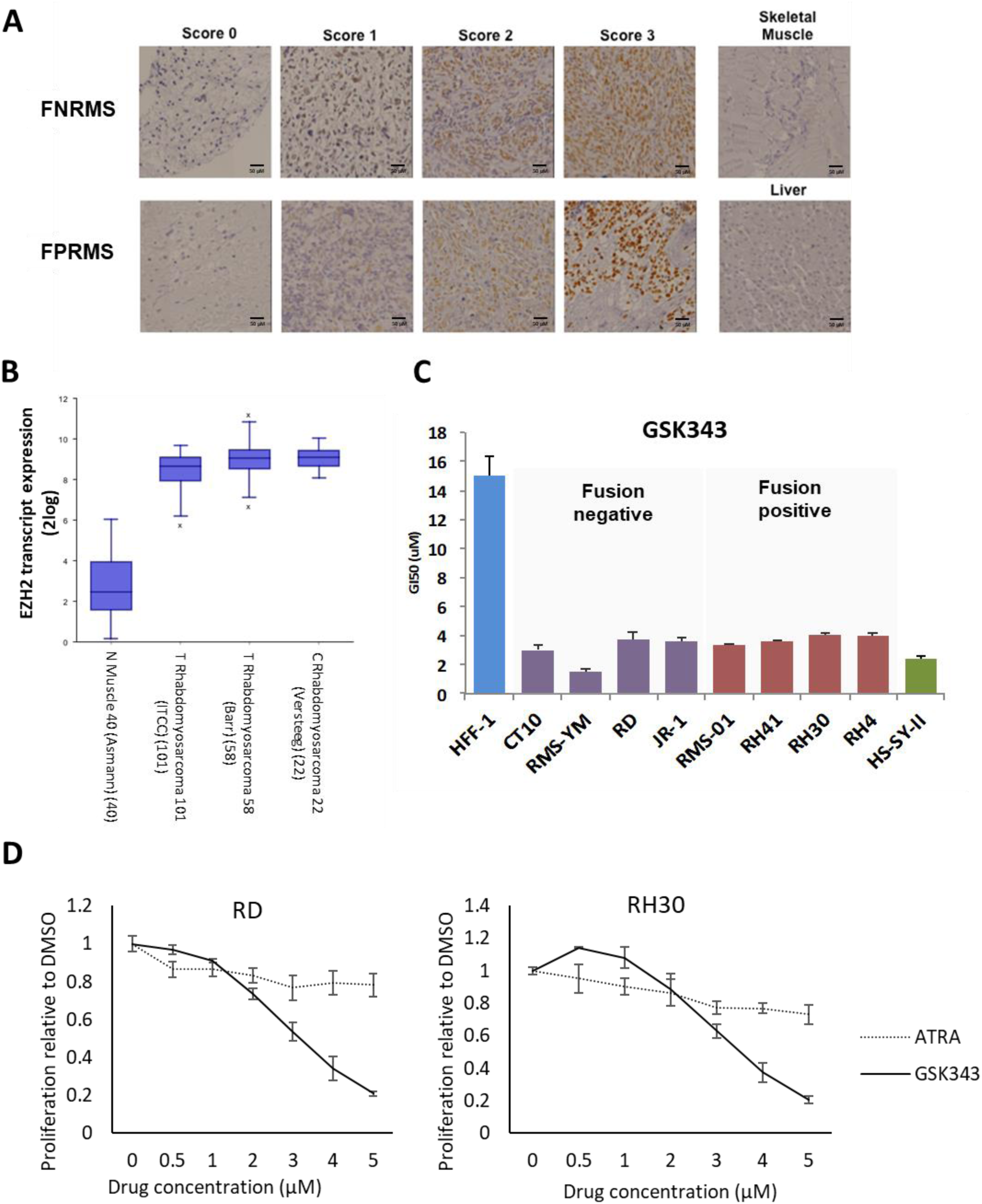
EZH2 is overexpressed in RMS and inhibition leads to a decrease in cell proliferation. **(A)** Photomicrograph of tissue microarray cores, showing examples of EZH2 protein expression in fusion negative RMS (FNRMS) and fusion positive RMS (FPRMS) samples scored as 0 (less than 5% positively stained cells), 1(weak), 2 (moderate), 3 (high). Scale bars = 50 μm. **(B)** EZH2 transcript is overexpressed in RMS tissue samples and cell lines. **(C)** GI50 after GSK343 treatment for RMS cell lines, HFF-1, and HS-SY-II. **(D)** Cell proliferation of RD and RH30 treated with indicated concentrations of GSK343 or ATRA for 6 days

**Table 1.** EZH2 protein staining in RMS patient samples by histology and clinical features**. FN** = Fusion Negative, FP = Fusion positive, U = Fusion status unknown

| Score | FN | FP | U | Total |
| --- | --- | --- | --- | --- |
| 0 | 37 | 9 | 12 | 58 |
| 1 | 85 | 31 | 9 | 125 |
| 2 | 61 | 16 | 5 | 82 |
| 3 | 17 | 0 | 0 | 17 |
| Total | 200 | 56 | 26 | 282 |

| Metastasis |  |  |  |
| --- | --- | --- | --- |
|  | 0 | 1 | Total |
| Neg | 48 | 8 | 56 |
| Pos | 174 | 42 | 216 |
| Total | 222 | 50 | 272 |

### 3.2. EZH2 inhibition reduces RMS cell proliferation

As the overexpression of EZH2 has been reported to support survival and proliferation in RMS cells [7], we tested the EZH2 inhibitor (EZH2i), GSK343 against a panel of RMS cell lines. These were compared with a normal fibroblast cell line, HFF-1 which are non-responsive to EZH2i, and the EZH2i-sensitive synovial sarcoma cell line, HS-SY-II [38]. Treatment of RMS cell lines with GSK343 lead to a significant decrease in cell proliferation and significantly lower GI50 over 6 days compared to control HFF1 cells (Figure 1C). Similar results were observed in all cell lines when tested with other EZH2i, GSK126 and UNC1999 (Figure S1, S2A). In contrast, EPZ6438 did not induce any significant changes in all RMS cell lines tested (Figure S3). RD and RH30 were selected for further work as representatives of FNRMS and FPRMS respectively. Additionally, we tested the inactive analog for UNC1999, UNC2400 [39] to check for specificity of EZH2 inhibition and found no significant changes in cell viability (Figure S2B).

We next compared the effects of GSK343 and differentiating agent, ATRA on cell proliferation. In contrast to GSK343, ATRA treatment showed a minimal decrease in proliferation in both cell lines (Figure 1D).

### 3.3. EZH2 inhibition potentiates ATRA treatment in FNRMS and FPRMS cell lines

As both EZH2 and retinoic acid signalling are known to influence differentiation, we sought to determine whether combination treatment EZH2i with RA treatment might be synergistic. Combination treatment resulted in a decrease in proliferation in both RD (Figure 2A) and RH30 (Figure 2B) cells at 6 and 10 days compared with the DMSO control (Figure 2B) and single agent treatment (Figure S4A-D). Evaluation of combination effect by EOB score indicated strong synergistic effects in both RD (Figure S5A) and RH30 (Figure S5B) by day 10.

**Figure 2.**
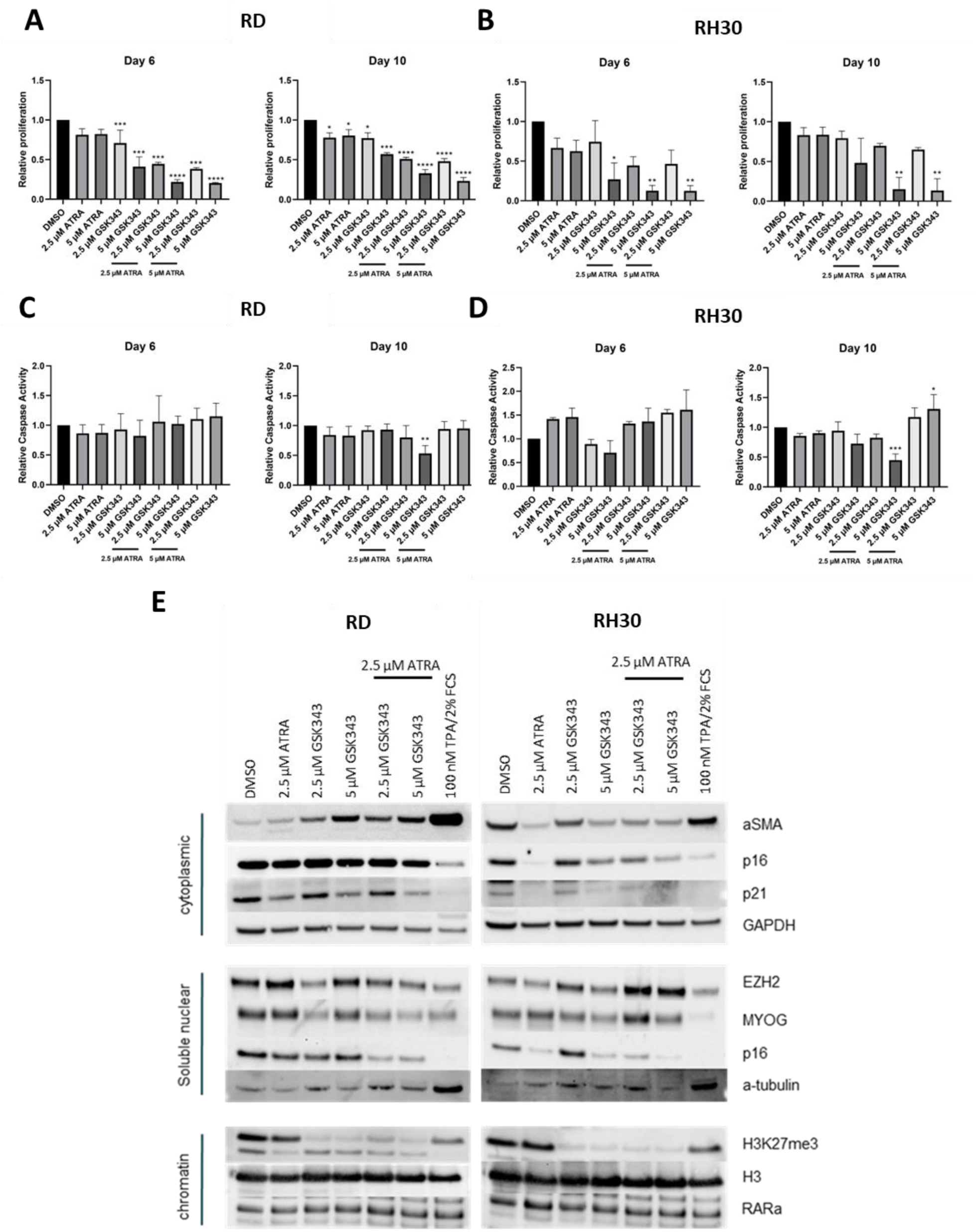
EZH2 inhibition potentiates the effect of ATRA in 2D. Cyquant proliferation assay of **(A)** RD and **(B)** RH30 after treatment with ATRA, GSK343 or in combination relative to DMSO control. Caspase signaling intensity of ATRA and GSK343 combination treatment in **(C)** RD and **(D)** RH30 cells relative to DMSO control. **(E)** Western blots of myogenic marker expression for fractionated RD and RH30 respectively treated with ATRA, GSK343, or combination for 6 days. Data shown are mean values from at least 2 independent experiments; bars, SD. One Way ANOVA was used to analyze statistical significance compared to the DMSO control (* p < 0.05, ** p < 0.01, *** p < 0.001, **** p < 0.001

### 3.4. Combination treatment induces differentiation in 2D FNRMS cells and apoptosis in FPRMS cells

As EZH2 inhibition has been linked to differentiation in FNRMS lines [13], versus apoptosis in FPRMS lines [7], we sought to determine whether the combination therapy was associated with a greater induction of myogenic differentiation and apoptosis respectively. Caspase activity levels did not show significant changes in RD after 6 days compared to the control and single agent (Figure 2C, S4E & F). At Day 10, an increase in caspase activity was observed at the combination treatments with 5 µM ATRA in the RH30 cells compared with control and signal agent (Figure 2D). These results suggests that the combination with 5 µM ATRA may induce an apoptotic phenotype in FPRMS.

Both cell lines were tested for the expression of myogenic markers, α-skeletal muscle actin (α-SMA) and myogenin (MYOG). Both RD and RH30, expressed MYOG in the absence of treatment (Figure 2E). Single agent treatments had little effect on MYOG expression, however combination treatment lead to a notable decrease in nuclear MYOG. The expression of α-SMA was noticeably different with RD showing no expression in the absence of drugs and a marked increase in response to 5 µM GSK343. This effect appears to be ameliorated in the combination treatment. Conversely, RH30 cells appear to express α-SMA constitutively with little variation due to either agent used alone and a marked reduction of detectable α-SMA with the combination treatment. These suggest that the combination therapy affects expression of regulators of myogenic differentiation in both RD and RH30, but the differences appear subtype dependent.

To determine if differentiation was the result of reduced cell proliferation, cell cycle proteins p16 (CDKN2A) and p21 (CDKN1A) were compared to cells differentiated using 12-O-tetradecanoylphorbol-13-acetate (TPA) (Figure 2E). The increased α-SMA and α-tubulin along with reduced expression of p21 and MYOG in RD cells implies that differentiation confers the growth inhibitory effects of the combination treatment. A reduction in the expression of cell cycle proteins in RH30 cells was also observed in the combination treatment but unlikely to be a consequence of differentiation as α-SMA was reduced in this line. This along with the results from the caspase assay implies that the growth inhibitory effect of the combination treatment in RH30 may be due to induction of the apoptosis pathway.

### 3.5. EZH2 inhibition potentiates ATRA treatment in FNRMS and FPRMS tumor spheroids

ATRA plus GSK343 treatment results in a reduction in cell growth in both FNRMS and FPRMS cells using 2D culture, so we aimed to determine whether these effects could be replicated in 3D spheroid models. Spheroids better represent *in vivo* tumor characteristics [40] and allow for longer term experiments important for observing epigenetic changes. Combination treatment induced a significant increase in RD cell viability at Day 14 with 5 µM GSK343 compared to the control and single agent (Figure 3A, S6A & B). In RH30 spheroids, a significant decrease in cell viability was observed at the combination treatment with 5 µM GSK343 and with combinations of both concentration of ATRA at Day 6. At Day 14, all treatments induced a significant decrease in cell viability, but combination treatments showed a stronger decrease compared to the control and single agent treatment (Figure 3B, S6C & D). These trends were also observed with spheroid volumes in the respective cell lines (Figure 3C-E, S6E-H), where combination treatment in FPRMS lines resulted in significant morphological changes consistent with apoptosis (Figure 3E, S6H).

**Figure 3.**
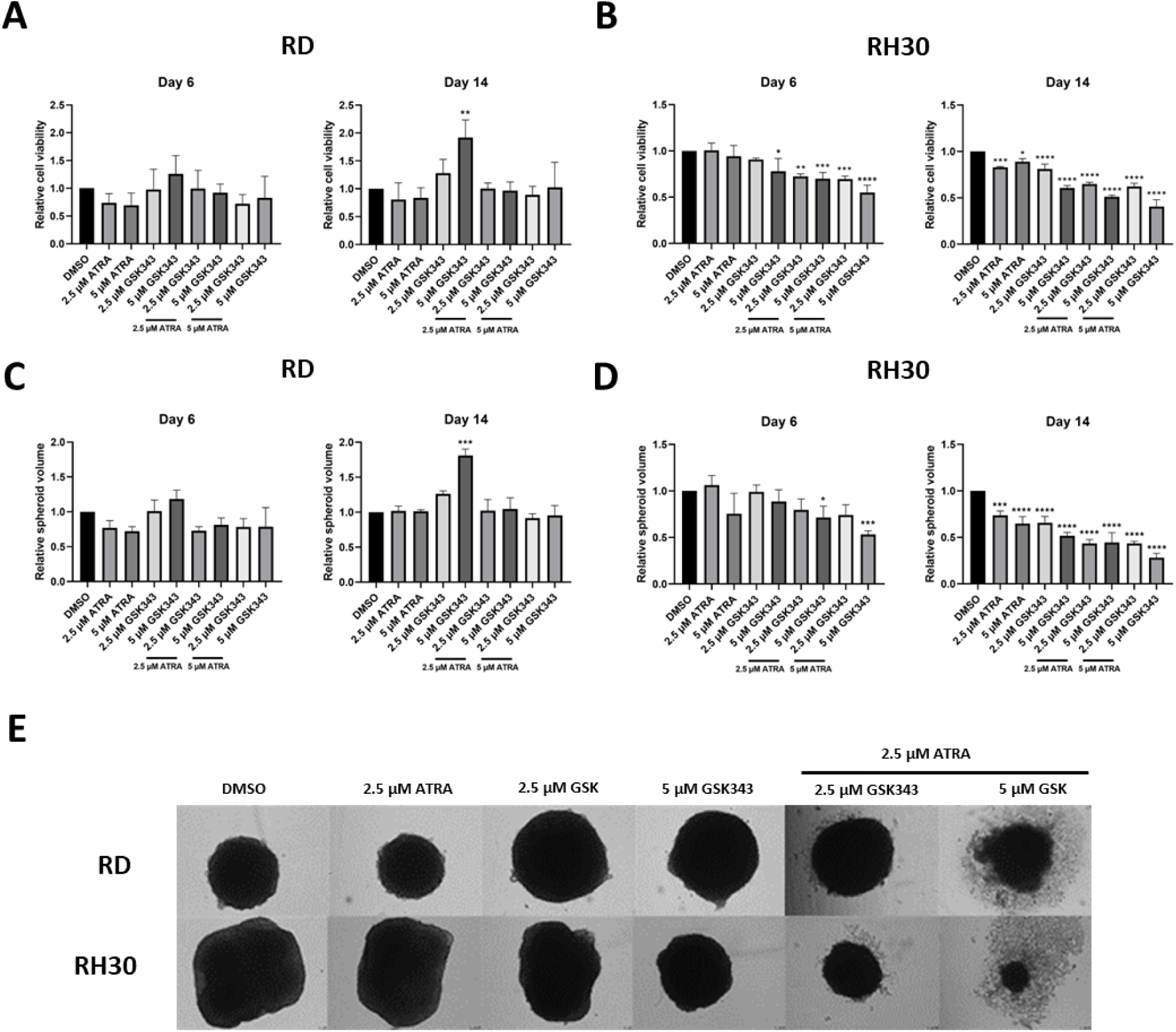
EZH2 inhibition potentiates the effect of ATRA in 3D RMS spheroids. 3D cell viability assay of **(A)** RD and **(B)** RH30 after treatment with ATRA, GSK343 or in combination at 6 and 14 days relative to DMSO control. **(C)** RD and **(D)** RH30 spheroid diameter relative to DMSO control after 6- and 14-days treatment Data shown are mean values from 3 independent experiments; bars, SD. One Way ANOVA was used to analyse statistical significance compared to the DMSO control (* p < 0.05, ** p < 0.01, *** p < 0.001, **** p < 0.001). **(E)** Microscope images of RD and RH30 spheroids treated with ATRA, GSK343, or combination for 14 days.

Combination treatment in RD spheroids showed strong synergy at Day 14 with ATRA and 5 µM GSK343 (Figure S7A). Combination treatment in RH30 spheroids showed strong synergy at Day 6 but this effect was reduced at Day 14 (Figure S7B). These results indicate that the inhibiting EZH2 and inducing RA signaling in FNRMS and FPRMS cells potentiated the growth inhibitory effects of the treatments.

### 3.6. Combination treatment induces differentiation in FNRMS spheroids and apoptosis in FPRMS spheroids

As 2D cells showed evidence of the upregulation of differentiation markers in RD and caspase activity in RH30, we determined whether the same could be seen in 3D spheroids. With combination treatment, RMS spheroids apoptose and disintegrate by Day 14 (Figure 3E) therefore further experiments were performed at Day 6. Doses of ATRA above 2.5 µM did not show an additional effect on viability and volume of spheroids therefore this concentration was used for further combination experiments.

H & E staining showed that RD spheroids treated with 5 µM GSK343, with or without 2.5 µM ATRA showed >60% differentiation characterized by features such as smaller nuclei, larger cytoplasm – changes in nuclei to cytoplasm ratio. This may explain the increasing spheroid size observed in Figure 3C and 3E (Figure 4A; Table S1). RH30 spheroids showed little evidence of differentiation (Figure 4B). The combination of GSK343 and ATRA showed anti-proliferative effects in RMS spheroids as evidenced by the reduction of the proliferative marker, Ki67 when compared to the DMSO control (Figure 4C). EZH2 inhibition was also observed as indicated by the decrease in H3K27me3 staining in both RMS spheroids treated with GSK343 alone and in combination (Figure 4D). The combination treatment showed evidence of myogenic differentiation in RD spheroids as seen by the presence of the terminal differentiation marker MHC (Figure 3H) and decrease in MYOG staining (Figure 4F). Conversely, there was little evidence of differentiation in RH30 spheroids by H&E staining, or IHC for the differentiation markers MHC and MYOG. Expression of the apoptotic marker, cleaved caspase 3 was positive in 5% of cells, localized to the cytoplasm and nucleus in GSK343 treated RH30 spheroids. However, no significant changes in cleaved caspase 3 were observed in the combination treatment (Figure 4G). Day 6 RH30 spheroids may be too early for induction of the apoptosis pathway. These data coupled with MTCS volume data from later timepoint suggest that combination treatment induced a pro-differentiation phenotype in RD spheroids versus likely cell death in RH30.

**Figure 4.**
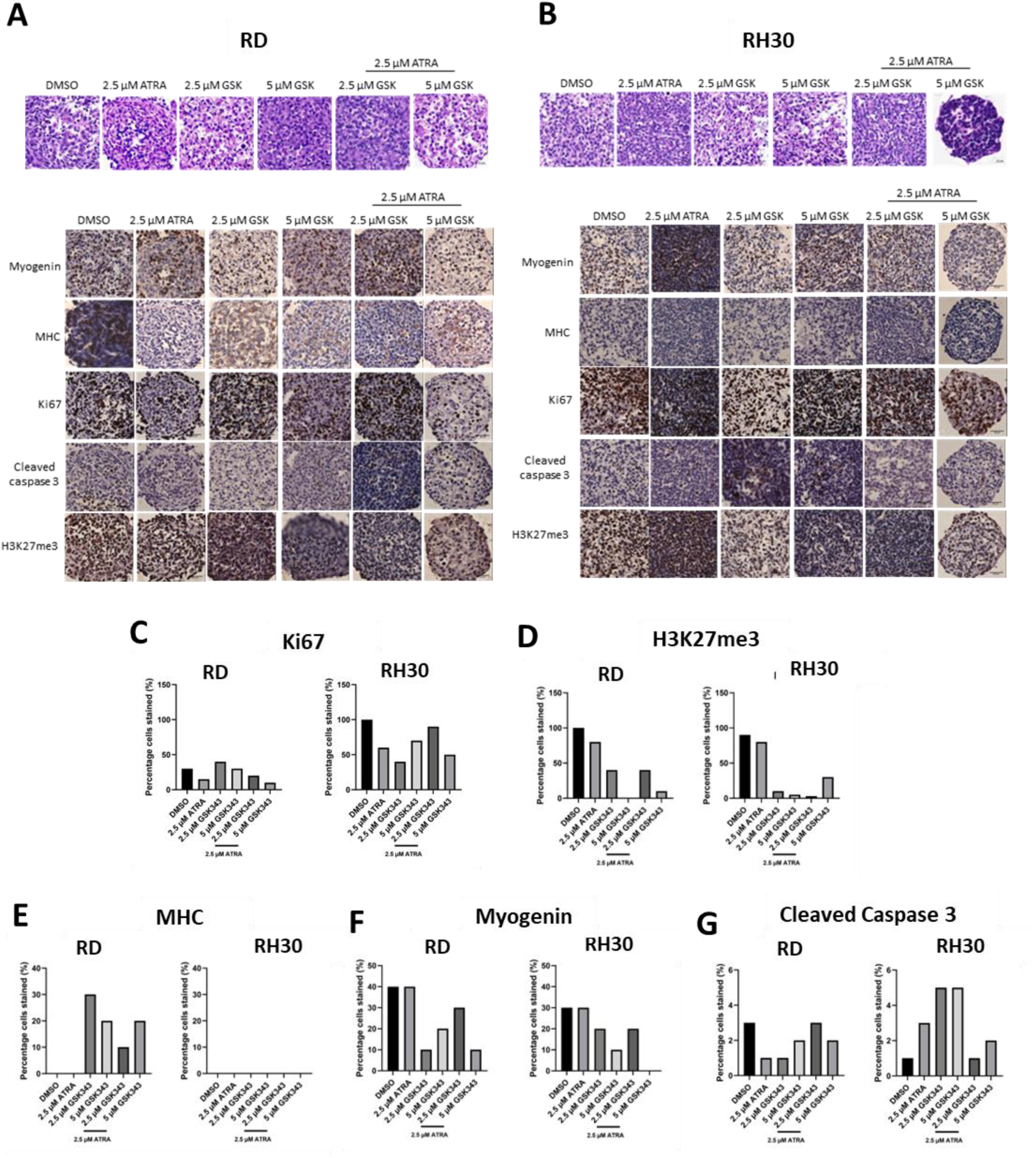
Combination treatment induces a pro-differentiation phenotype in RD spheroids and cell death in RH30 spheroids. Representative images of hematoxylin and eosin staining (H & E) and immunohistochemistry (IHC) of proteins in RD **(A)** and RH30 **(B)** spheroids after 6 days of treatment. Percentage of cells stained in immunohistochemistry (IHC) of proteins in RD and RH30 spheroids after 6 days treatment using antibodies targeting **(C)** Ki67, **(D)** H3K27me3, **(E)** Myosin Heavy Chain (MHC), **(F)** Myogenin and **(G)** Cleaved Caspase 3.

We next sought to identify pathways and genes that are differentially expressed between treatments to further understand the mechanism behind combination treatment and the different phenotypes observed in the FPRMS and FNRMS cell lines.

### 3.7. ATRA treatment alone induced an upregulation of genes involved in the interferon response pathway in FPRMS cells

As ATRA can initiate RA signaling through the binding of RARs, we identified genes bound and regulated by RARα through the integration of ChIP-seq and RNA-seq data to understand the mechanism of action. RNA-seq was performed on RMS cells treated with 2.5 µM ATRA for 6 days. In RD, 100 genes were significantly upregulated compared to DMSO control, while 14 genes were downregulated (FC > 2, adj p value < 0.05; Figure S8A), with significant upregulation of differentially expressed genes (DEGs) associated with estrogen early response and KRAS signalling by gene set enrichment analysis (GSEA) (Figure S8B). In RH30, 339 genes were upregulated, compared to DMSO control, and 135 genes were downregulated (FC > 2, adj p value < 0.05; Figure S8C), with significant upregulation of DEGs associated with the immune response Figure S8D).

Integration of ChIP-seq and RNA-seq data showed the low degree of overlap in RARα binding domains with changes in gene expression in both RMS cell which may be due to the timepoint used for RNA-seq as RAR-responsive genes have been shown to respond rapidly to RA-stimulation [41, 42] (Figure S8E & F).

### 3.8. GSK343 treatment alone induced an upregulation of genes involved in the myogenesis pathway in RD cells

To understand the underlying mechanism of EZH2 inhibition, RNA-seq was performed on RMS cells treated with 5 µM GSK343 for 6 days and compared to DMSO control. In RD, 858 genes were significantly upregulated and 20 significantly downregulated (FC > 2, adj. p value < 0.05), with significant upregulation of DEGs associated with the canonical MSigDB myogenesis pathway (Figure S9A and B). In RH30 cells, 457 genes were significantly upregulated and 52 significantly downregulated (FC > 2, adj. P value < 0.05) with similar enrichment in pathways seen in RD (Figure S9C and D). Overall, EZH2 inhibition resulted in an upregulation of cholesterol homeostasis genes in both cells (RD - 25/74, RH30 - 10/74; Figure 6D) suggesting that GSK343 targets RMS cells on a metabolic level, in addition to myogenesis in FNRMS.

We next investigated EZH2 binding using ChIP-seq on untreated RMS cells. Integration of the ChIP-seq and RNA-seq data showed 8 genes that were downregulated, and 414 genes were upregulated in RD that showed an overlap with EZH2 peaks (Figure S9E). In RH30 cells, 44 genes were downregulated whilst 418 genes were upregulated that EZH2 may bind to (Figure S9F). Pathway analysis using MSigDB canonical pathway gene sets and the DEGs that are potentially regulated by EZH2 indicates that EZH2 may play a role in regulating cholesterol homeostasis and myogenesis in both RMS subtypes (Figure S9G). Using a lower concentration of GSK343 (2.5 µM) showed an upregulation in genes involved in the same pathways however to a lesser degree compared to the higher concentration (5 µM) (Figure S10).

Epigenetic reprogramming of H3K27me3 marks by EZH2 in RMS cells was also explored using ChIP-seq. We found that there was a large overlap between EZH2 and H3K27me3 peaks in both RMS subtypes which correlates with the role of EZH2 in the trimethylation of H3K27 (Figure S11).

### 3.9. Combination treatment strongly induces an additive upregulation in myogenesis in FNRMS and the interferon response pathway in FPRMS

To understand the underlying mechanism of the potentiating effect observed in the combination treatment, RNA-seq was performed on RMS cells treated with 2.5 µM ATRA and 5 µM GSK343 for 6 days. More DEGs were seen in RD cells after combination treatment (versus control) compared to single agent GSK343(combination – 1275; GSK343 – 878), with significant enrichment in the myogenesis pathway by GSEA (Figure 5A-C) compared to GSK343 alone (combination - 42/200, GSK343 - 45). Similarly, after combination treatment in RH30, an increase in significant DEGs was observed compared to single agent ATRA (combination – 1651; ATRA – 474), with significant enrichment in the IFN-α pathway and in other immune response canonical pathways (Figure 5D and E). Treatment of RMS cells with 2.5 µM ATRA/2.5 µM GSK343 showed a similar trend in the upregulation of the same pathways as treatment with 2.5 µM ATRA/5 µM GSK343 (Figure S11, S12A-C).

**Figure 5.**
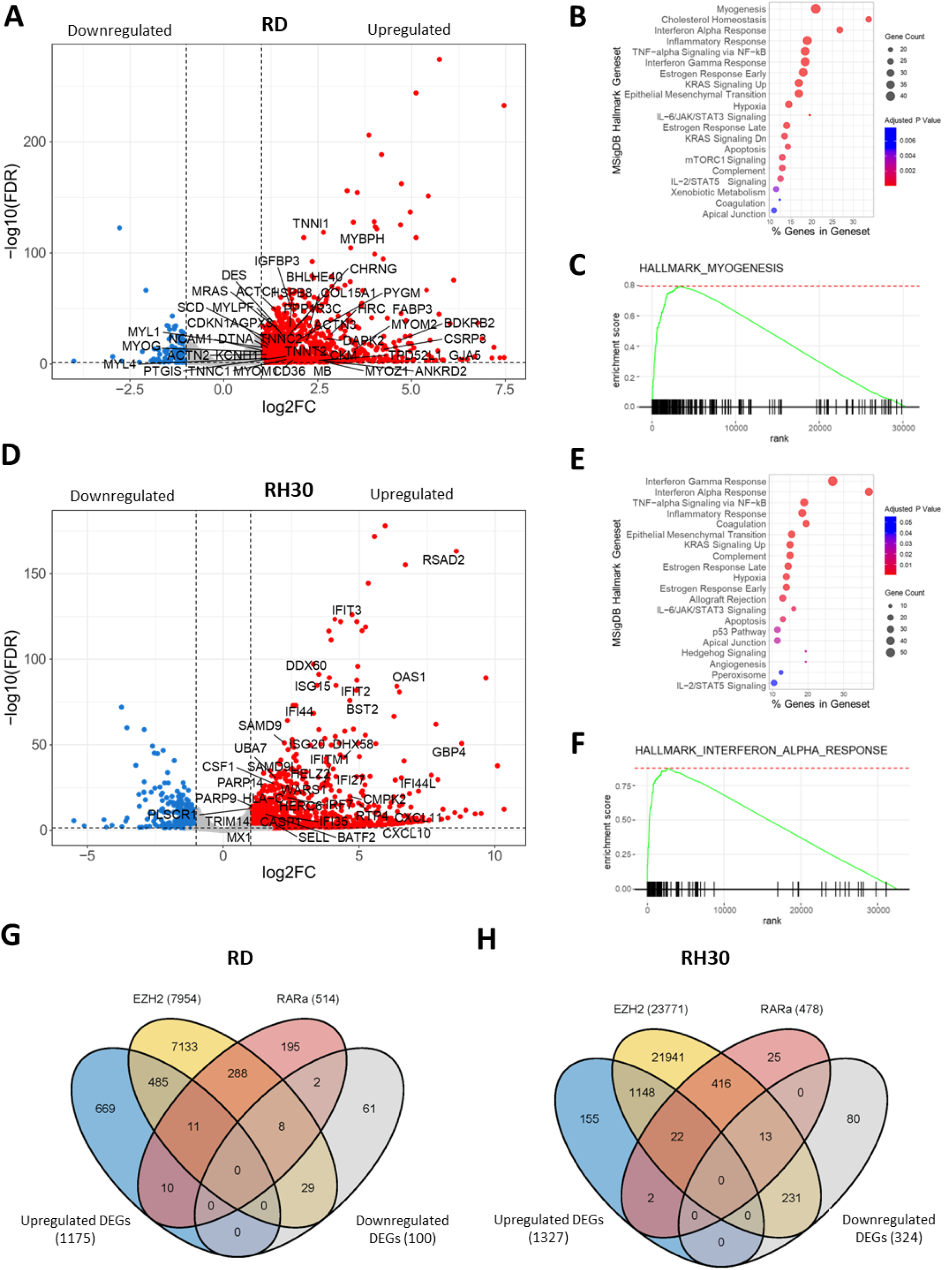
Integration of RNA-seq and ChIP-seq data in RMS cells treated with a combination of 2.5 µM ATRA and 5 µM GSK343 compared to DMSO control. **(A)** Volcano plot of genes involved in myogenesis in RD cells after treatment. **(B)** Gene ontology analysis of processes that are differentially regulated in RD cells treated with combination of 2.5 µM ATRA and 5 µM GSK343. **(C)** GSEA shows positive enrichment of myogenesis genes in RD after combination treatment. **(D)** Volcano plot of genes involved in the interferon α response pathway in RH30 cells after treatment. **(E)** Gene ontology analysis of processes that are differentially regulated in RH30 cells treated with combination of 2.5 µM ATRA and 5 µM GSK343. **(F)** GSEA shows positive enrichment of genes involved in the interferon α response in RH30 after combination treatment. Venn diagram showing the overlap of number of DEGs with EZH2 and RARα peaks identified from ChIP-seq in **(G)** RD and **(H)** RH30 cells.

Overlap of RARα and EZH2 peaks were analyzed to investigate whether they were present in the same genes, indicating whether they might regulate overlapping targets. There was a significant overlap in genes located near peaks for EZH2 and for RARa in both RD and RH30; in RD, RARα and EZH2 peaks were both observed in 307 genes (One-sided Fisher’s exact test, p < 0.0001; Figure S13A) and in RH30, there was an overlap of 451 genes (One-sided Fisher’s exact test, p < 0.0001; Figure S13C). Pathway analysis revealed that RARα and EZH2 bind to and regulate the same genes in TNF-α signaling via NFkB in both cell lines in addition to TGFβ signaling in RH30 (Figure S13B and D). Under all treatments tested, negatively enriched canonical pathways were cell cycle related (E2F targets, MYC targets, and G2M checkpoint) and/or oxidative phosphorylation in both subtypes of RMS (Figure S14 and S15).

To investigate if RARα and EZH2 regulated the same genes in the combination treatment, ChIP-seq was used to assay RARα and EZH2 binding sites in untreated RMS cells. RARα and EZH2 peaks were detected in 11 of the upregulated genes and 8 of the downregulated genes in the RD cell differential gene expression analysis (Figure 5G). Specifically, *CCND2* is upregulated in the combination treatment and has a role in cell cycle and differentiation [43]. In RH30 cells, RARα and EZH2 peaks were observed in 22 upregulated genes, including immunoregulatory gene *VSIR*, and 13 downregulated genes from the differential gene expression analysis (Figure 5H). Few RARα peaks were present in DEGs in both RMS cell lines perhaps owing to the single timepoint used for RNA-seq, however EZH2 peaks were detected in 496 upregulated DEGS and 37 downregulated DEGs in RD cells (Figure S16D). In RH30 cells, EZH2 peaks overlapped with 1142 upregulated DEGs and 235 downregulated DEGs (Figure S16E). Pathway analysis on integrated ChIP-seq and RNA-seq data showed that EZH2 bound to genes involved in myogenesis and inflammatory response including the interferon response (Figure S16F). Collectively, these observations suggest that combination EZH2i and ATRA treatment may be an effective therapy in both FN and FP RMS by enhancing myogenesis and inflammation, respectively, further suggesting that EZH2i/ATRA treated FPRMS tumour cells may respond to immune-based therapies.

## 4. Discussion

Differentiation therapy shows promise in the treatment of cancer, showing positive outcomes in certain cancer types [44]. The results of this study provide evidence suggesting the use of EZH2i and RA treatment can induce myogenic differentiation, inhibit proliferation, and increase apoptosis as potential treatment for RMS patients. In all agents tested in this study, Myc targets and oxidative phosphorylation were downregulated in RMS which suggests in general, that the anti-proliferative effect of these treatments may also involve the targeting of cancer metabolism and proliferation.

Retinoic acid receptors, activated by RA ligands, act as a transcription factor to enhance the expression of specific myogenic genes [45] and RMS xenografts treated with ATRA showed enhanced MHC and decreased MYOG indicative of terminal muscle cell differentiation [21]. Contrastingly, several RMS cell lines, including RD and RH30 show limited response to RA as indicated by lack of reduction in cell growth or induction of myogenic differentiation, which was previously suggested to be due to low RAR expression [46]. Consistent with this, our results also show that ATRA treatment alone in both RD and RH30 cell lines had little effect on growth and differentiation. However, this is unlikely due to RAR expression as RARα is expressed in our cell lines. As PRC2-EZH2 has been shown to be involved in suppressing the RA signaling pathway [47–49], inhibiting EZH2 could allow RA ligand binding to facilitate signaling which may explain why ATRA alone was ineffective as a single agent treatment.

Our data indicate that the anti-cancer effect of EZH2 inhibition in RMS cells may partly be due to the dysregulation of cholesterol homeostasis as revealed by the RNA-seq and ChIP-seq data. Similar findings were observed in head and neck squamous carcinoma [50] and hepatocellular carcinoma [50] where EZH2i resulted in altered cholesterol synthesis. Dysregulation of cholesterol homeostasis can induce cell cycle arrest and apoptosis through the activation of specific transcription factors [51].

We show that combination treatment with EZH2i and RA in RMS 2D cell culture showed an anti-proliferative effect. Combination treatment resulted in a pro-differentiation phenotype in FNRMS versus a pro-apoptotic phenotype in the FPRMS in both 2D culture and 3D spheroid. Epigenetic profiling revealed that the pro-differentiation phenotype observed in FNRMS appeared to be driven by EZH2i, as combination treatment with ATRA resulted in synergism by upregulating more genes involved in myogenesis pathways. Upregulation of myogenic markers were also observed in 2D cells and spheroid in FNRMS which were not seen in FPRMS. This supports the role of oncogenic role of EZH2 in FNRMS via the suppression of myogenic differentiation. Our results show that RARα and EZH2 did not show many overlaps in DEGs of the treated RMS cells. This may be due to the timing of the experiment, the cells were harvested at Day 6 to capture the targets that are influenced by EZH2 inhibition (i.e., after changes in the H3K27me3 mark) however this time point may be too late to observe direct targets of RARα. Further investigation is required to understand what targets RARα regulate and if RARα regulates the same genes as EZH2 in the combination treatment.

In FPRMS, the pro-apoptosis phenotype appeared to be driven by ATRA potentiated by EZH2 inhibition. This pro-apoptotic phenotype in FPRMS has been observed in a number of previous studies where RH30 cells treated with EZH2i resulted in a dose-dependent increase in apoptosis rather than differentiation [13, 14]. This apoptotic phenotype may be immune-related as genes of the IFN-α pathway were upregulated. RA was reported to induce the secretion IFN-α in various human cell lines [52]. IFN-α has a reported role in inducing apoptosis in malignant cancer cells [53]. Whilst ATRA alone induced an upregulation of these pathways, combination with EZH2i was required to reduce viability and proliferation of FPRMS. Our data showed that EZH2 may regulate the genes involved in similar signaling pathways to those upregulated in ATRA treatment in FPRMS. Inhibition of EZH2 appeared to potentiate the effect of ATRA as more genes involved in the interferon α and γ response pathway were upregulated.

The difference in cellular response to EZH2 inhibition and/or RA signaling in the subtypes may be attributed to the different signaling pathways established due to the fusion oncogene. Comparison of primary RMS tumors and FNRMS transduced with PAX3-FOXO1 constructions revealed DEGs involved in apoptosis, cell death and negative regulation of cell proliferation [54]. Apoptosis appeared to be present in FPRMS tumors, however the baseline level was not sufficient to prevent tumor formation [55]. This highlights the paradoxical role of PAX3-FOXO1 where it can be oncogenic or anti- cancer. As the PAX3-FOXO1 fusion gene inhibits myogenic differentiation in FPRMS [56], its presence could determine why the FPRMS favors apoptosis rather than differentiation upon treatment with differentiating agents.

Overall, our findings provide insight into the mechanism that drives the anti-cancer effect of the EZH2/RA single agent and combination treatment, and the effect determined by the presence of the fusion oncogene. Ideally *in vivo* preclinical assessment of the combination would provide further rationale for the use of this combination in RMS, however the lack of appropriate immunocompetent models of RMS limits the ability to effectively test this. Nevertheless, our results support the potential use of this combination therapy for the treatment of both FNRMS and FPRMS.

## Financial Support

This work was funded by Children with Cancer UK [grant 14-177], Sarcoma UK [grant SUK206.2017] and The Schottlander Research Charitable Trust.

## CRediT authorship contribution statement

**Eleanor O’Brien:** Formal analysis, Investigation, Methodology, Validation, Writing – original draft. **Carmen Tse:** Formal analysis, Investigation, Methodology, Validation, Visualization, Writing – original draft. **Ian Tracy:** Formal analysis, Investigation, Methodology, Validation, Writing – original draft. **Ian Reddin:** Data curation, Formal analysis, Investigation, Methodology, Visualization, Writing – original draft. **Joanna Selfe:** Data curation, Formal analysis. **Jane Gibson:** Data curation, Formal analysis, Investigation. **William Tapper:** Data curation, Formal analysis, Investigation. **Reuben J Pengelly:** Data curation, Formal analysis, Investigation. **Jinhui Gao:** Formal analysis, Investigation. **Ewa Aladowicz:** Investigation, Formal analysis. **Gemma Peets:** Formal analysis. **Khin Thway:** Formal analysis. **Sergey Popov:** Formal analysis. **Anna Kelsey:** Formal analysis. **Timothy J Underwood:** Resources, Supervision, Writing – review & editing. **Janet Shipley:** Funding acquisition, Project administration, Resources, Software, Supervision, Writing – review & editing. **Zoë S Walters:** Conceptualization, Funding acquisition, Methodology, Project administration, Resources, Software, Supervision, Validation, Visualization, Writing – original draft.

## Declaration of competing Interest

All authors have no conflict of interest to declare.

## Acknowledgements

We would like to thank Dr Patrick Duriez for his technical assistance in the apoptosis assay.

## Ethics approval and consent to participate

Formalin-fixed paraffin-embedded (FFPE) samples from UK patients enrolled on the MMT89, MMT95 and MMT98 trials from the International Society of Pediatric Oncology were collected from multiple UK centers and used for immunohistochemistry analyses. The study was conducted according to the guidelines of the Declaration of Helsinki and approved by the Local Research Ethics Committee protocol 1836 and UK Multi-Regional Research Ethics Committee 98/4/023 (16/11/06).

## Abbreviations

RMS: Rhabdomyosarcoma
ERMS: embryonal RMS
ARMS: alveolar RMS
FNRMS: fusion negative RMS
FPRMS: fusion positive RMS
RA: retinoic acid
ATRA: All-*trans* retinoic acid
EZH2: Enhancer of Zeste Homolog 2
PRC2: Polycomb Repressive Complex 2
EZH2i: EZH2 inhibitor
TMS: Tissue Microarray
EOB: Excess Over Bliss Score
MYOG: Myogenin
RNA-seq: RNA- sequencing
ChIP: Chromatin Immunoprecipitation
ChIP-seq: ChIP-sequencing
DEGs: Differentially Expressed Genes
GSEA: Gene Set Enichment Analysis

---

**Figure S1.**
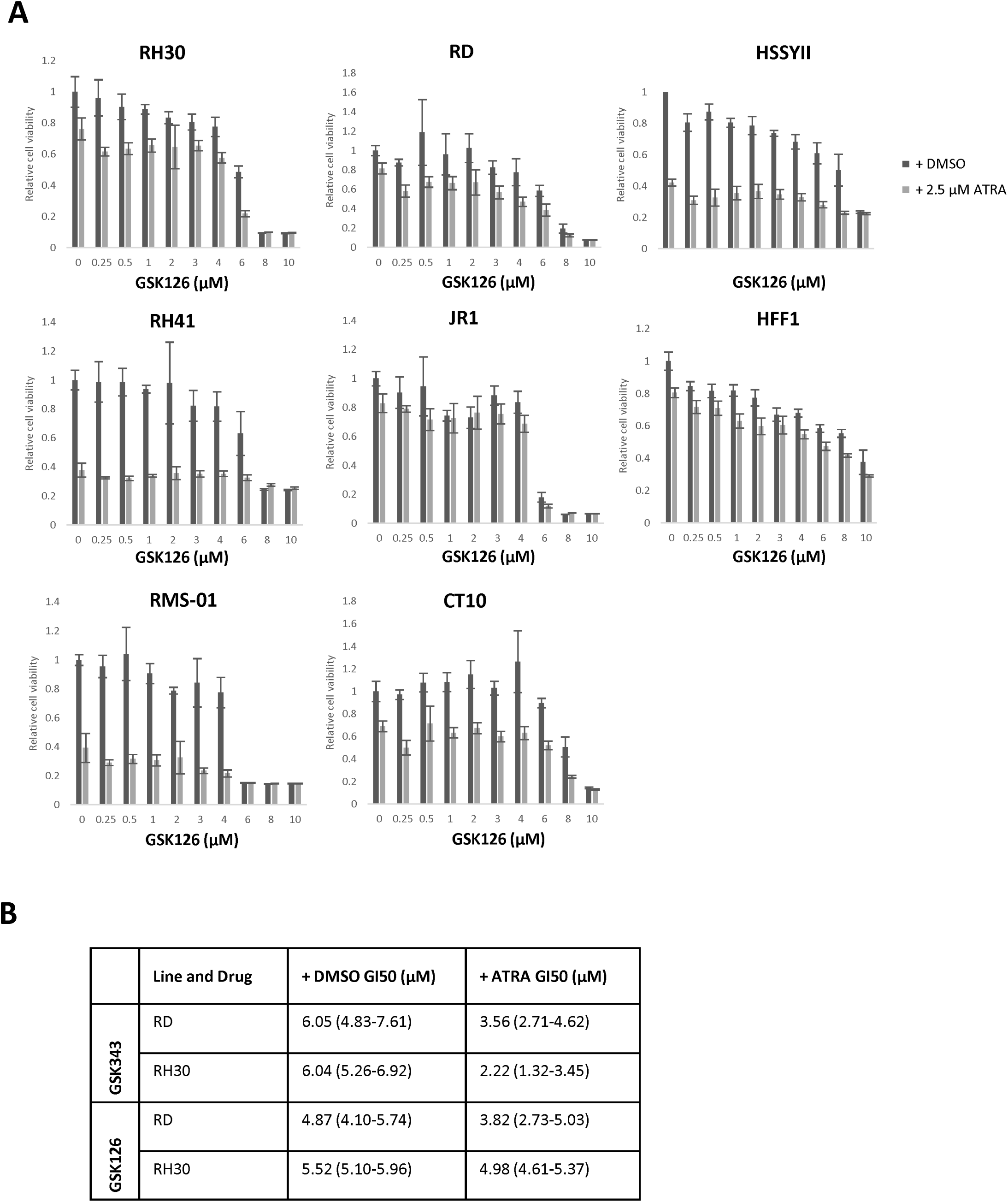
(A) Dose Response for the EZH2 inhibitor GSK126 in ARMS and ERMS cell lines combined with DMSO control or 2.5uM ATRA for 6 days. HSSYII and HFF1 used as a positive and negative control respectively. (B) GI50 with confidence intervals for GSK343 and GSK126 combined with 2.5uM ATRA.

**Figure S2.**
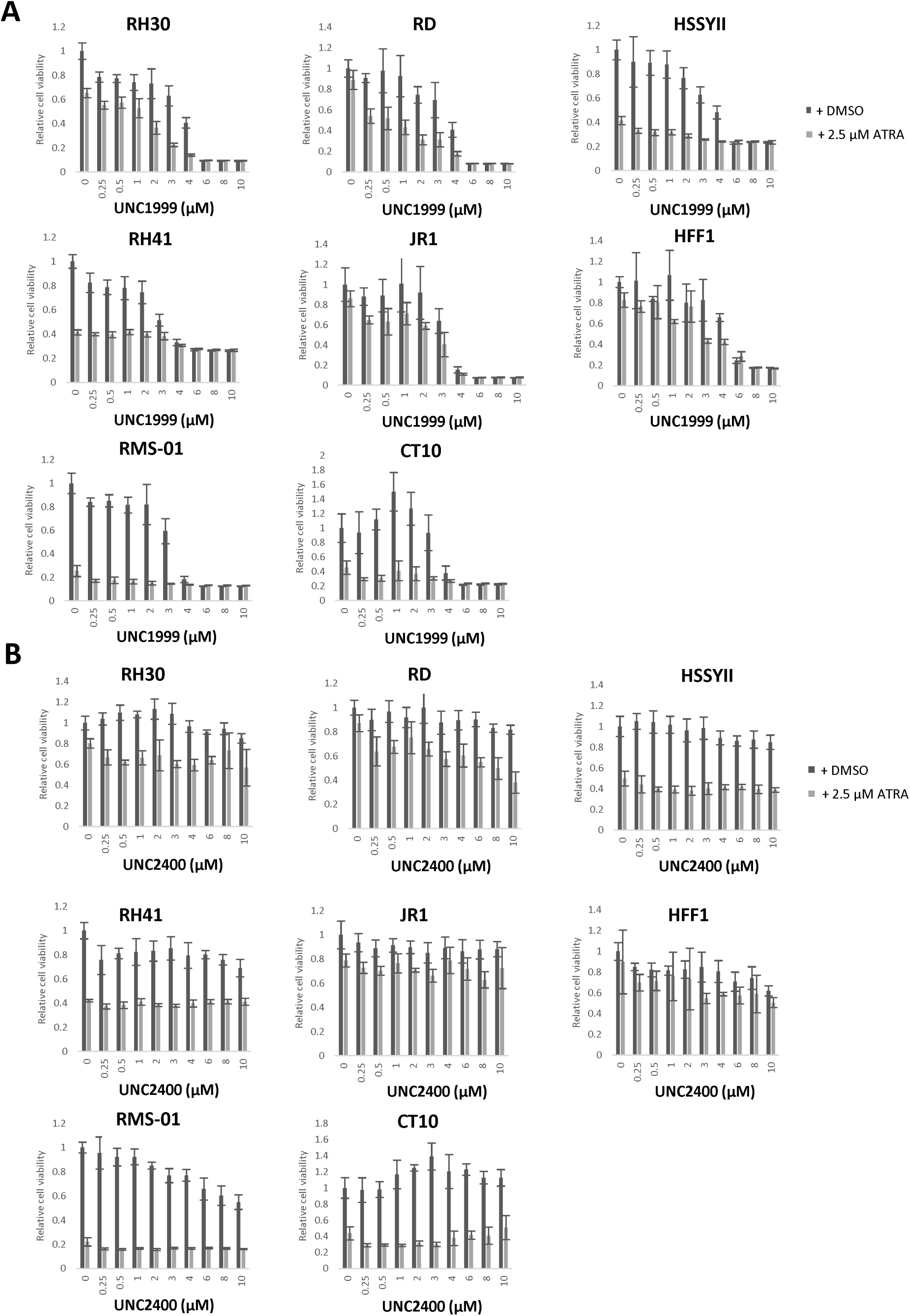
Dose Response for the EZH2 inhibitor UNC1999 **(A)** and the inactive analogue UNC2400 **(B)** in FP-RMS and FN-RMS cell lines combined with DMSO control or 2.5uM ATRA for 6 days. HSSYII and HFF1 used as a positive and negative controls respectively.

**Figure S3.**
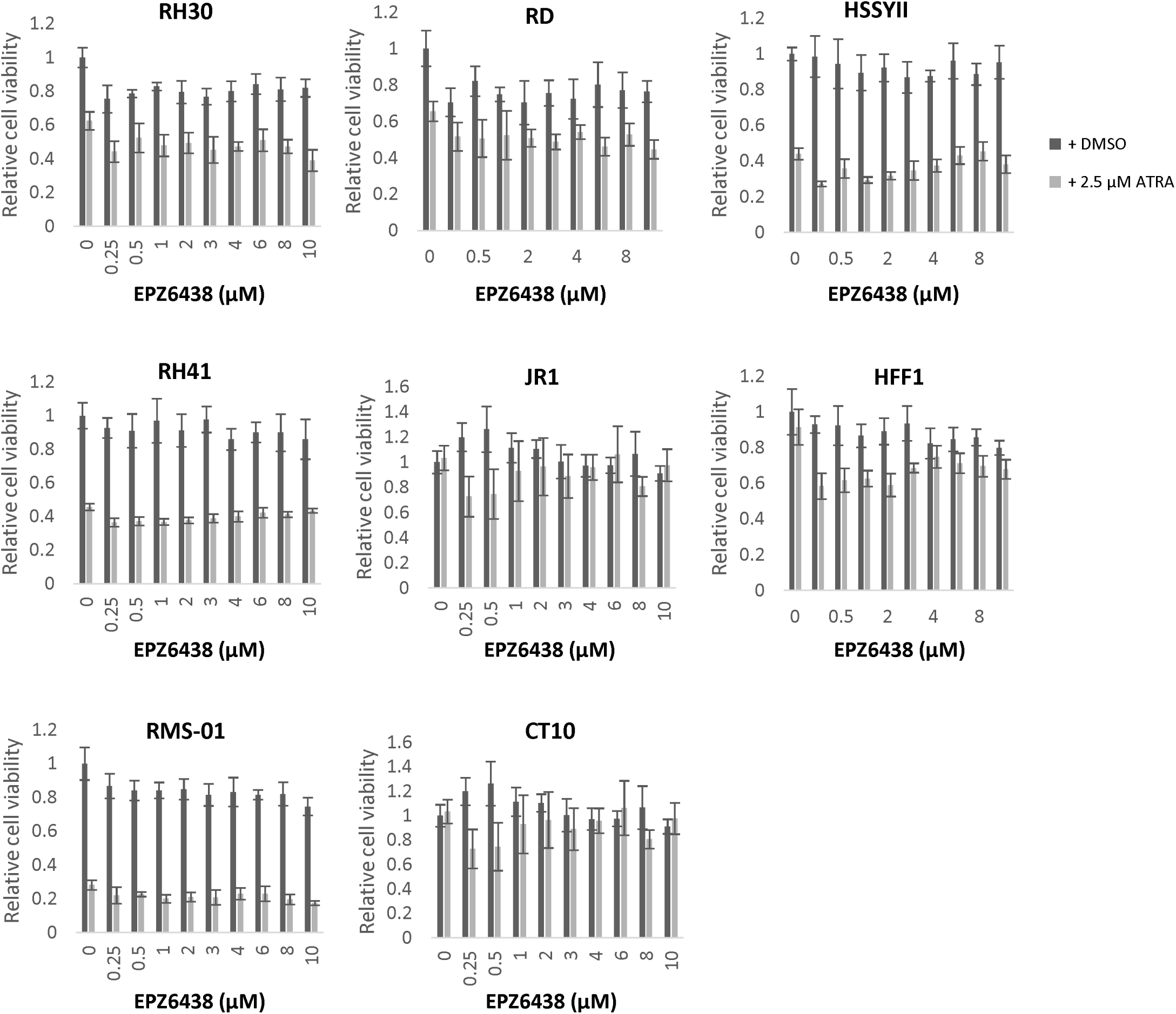
Dose Response for the EZH2 inhibitor EPZ6438 in FP-RMS and FN-RMS cell lines combined with DMSO control or 2.5 µM ATRA for 6 days. HSSYII and HFF1 used as a positive and negative control respectively

**Figure S4.**
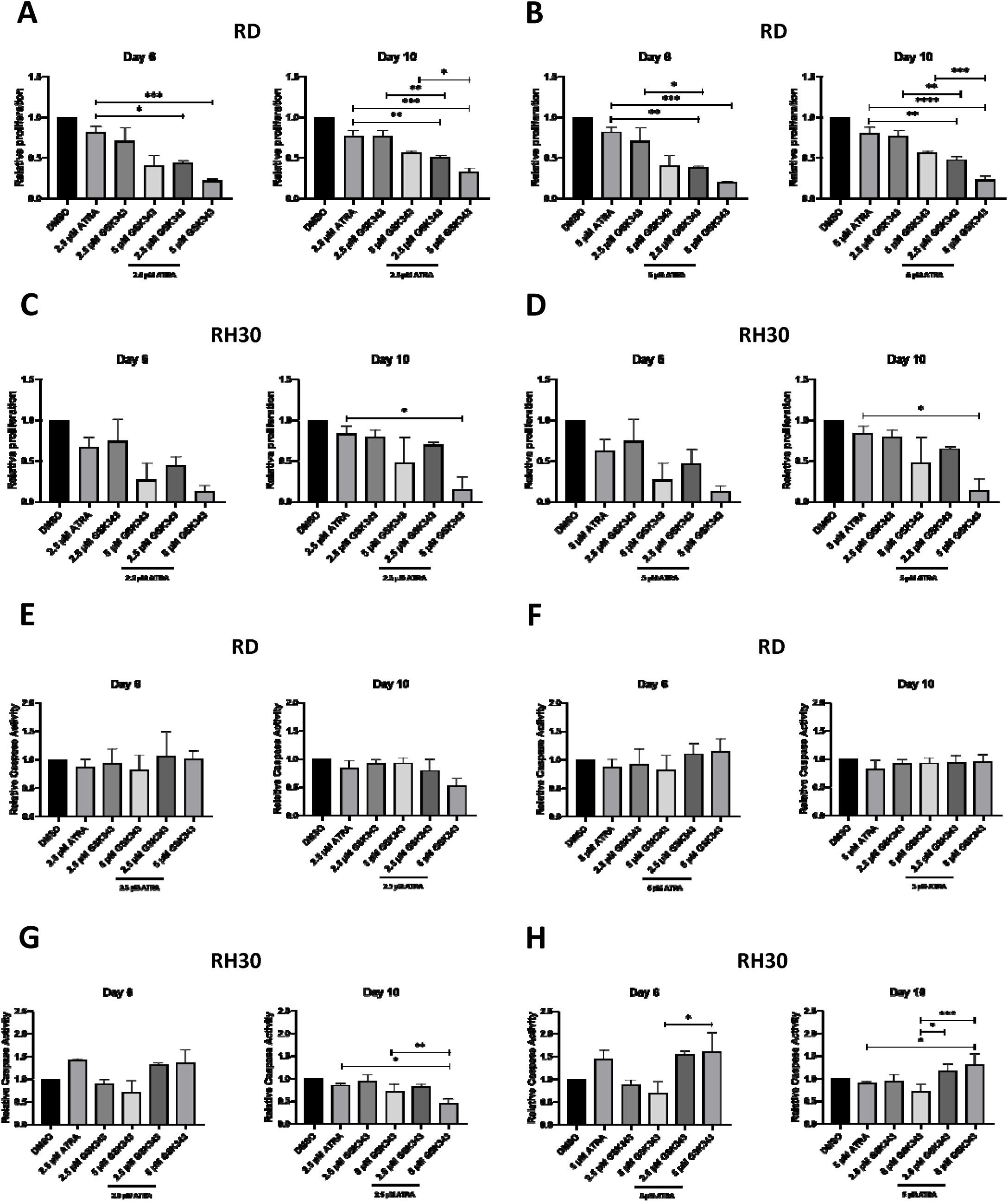
EZH2 inhibition potentiates effect of ATRA in 2D. Cyquant proliferation assay after treatment with 2.5 µM ATRA, GSK343 or in combination relative to DMSO control in RD (A) and RH30 (B) and treatment with 5 µM ATRA, GSK343 or in combination relative to DMSO control in RD (C) and RH30 (D). Caspase signaling intensity after treatment with 2.5 µM ATRA, GSK343 or in combination relative to DMSO control in RD (E) and RH30 (F) and treatment with 5 µM ATRA, GSK343 or in combination relative to DMSO control in RD (G) and RH30 (H). Data shown are mean values from at least 2 independent experiments; bars, SD. One Way ANOVA was used to analyse statistical significance compared to single agent (* p < 0.05, ** p < 0.01, *** p < 0.001, **** p < 0.001.

**Figures S5.**
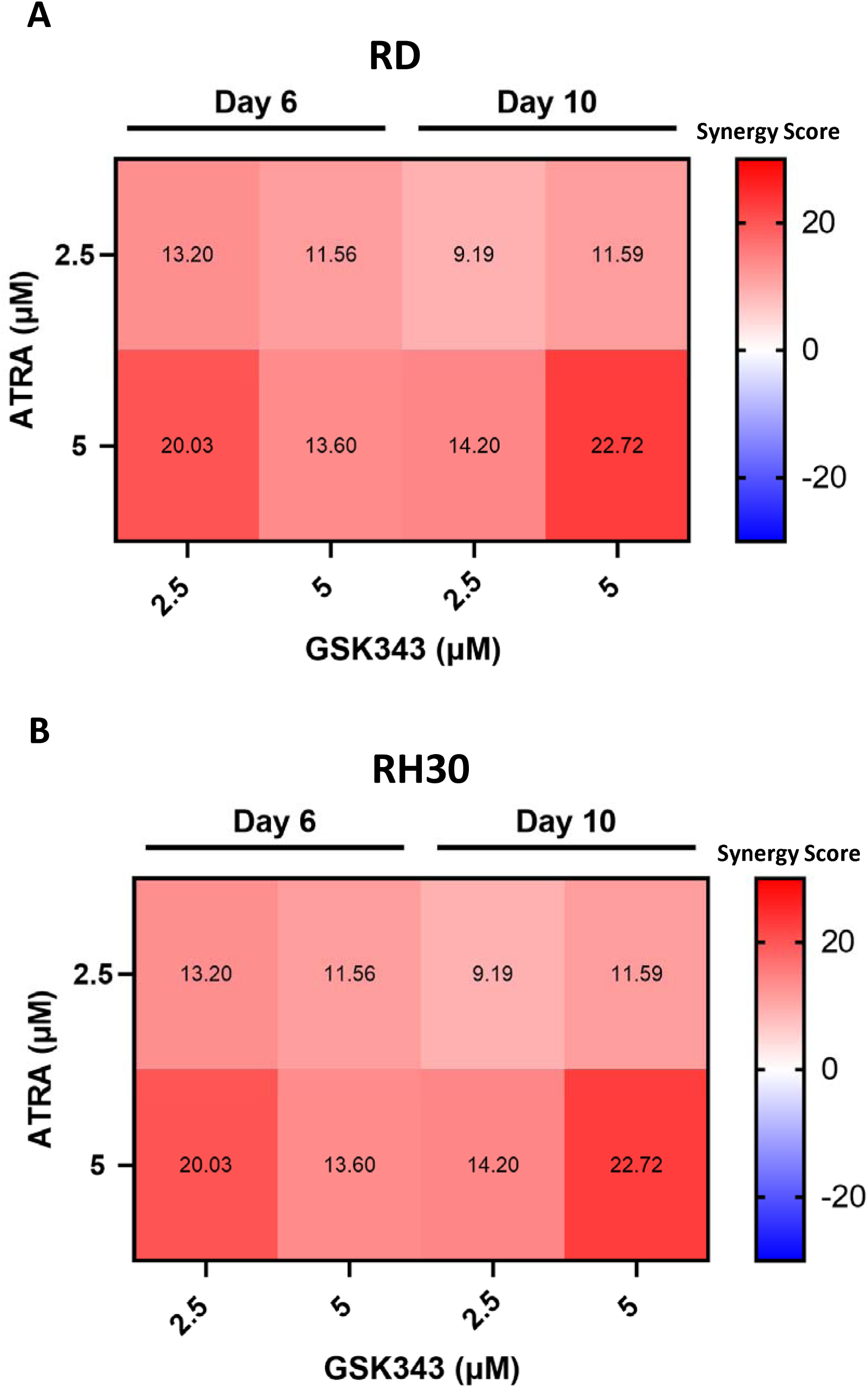
**Excess over Bliss calculations of 2D RD (A) and RH30 cells (B) treated with combinations of ATRA and GSK343.** The predicted Bliss response was subtracted from the experimentally observed inhibition. Scores that are > 0 represent synergy and < 0 represents antagonism.

**Figure S6.**
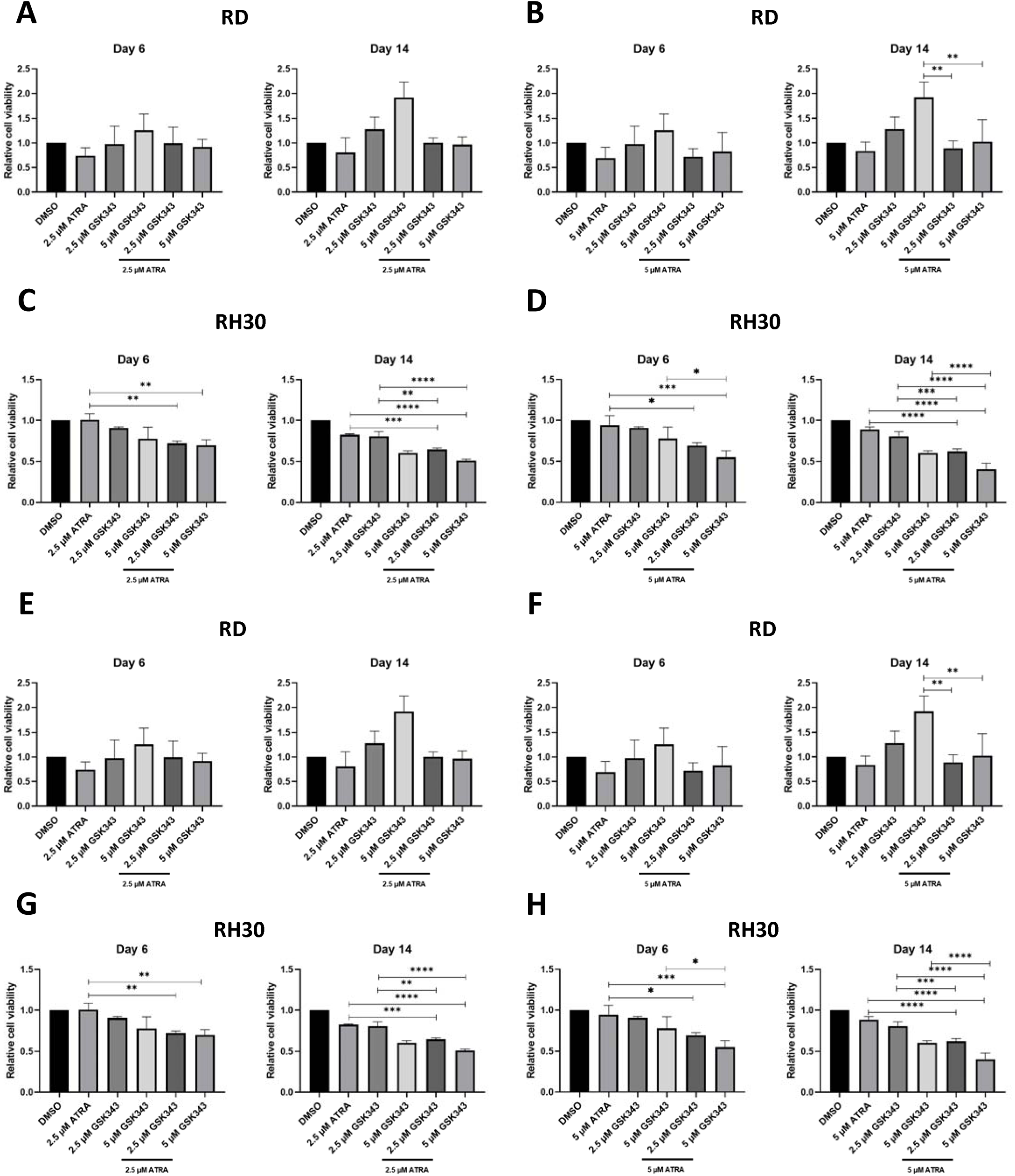
**EZH2 inhibition potentiates effect of ATRA in 3D.** Cyquant proliferation assay after treatment with 2.5 µM ATRA, GSK343 or in combination relative to DMSO control in RD **(A)** and RH30 **(B)** and treatment with 5 µM ATRA, GSK343 or in combination relative to DMSO control in RD **(C)** and RH30 **(D)**. Caspase signaling intensity after treatment with 2.5 µM ATRA, GSK343 or in combination relative to DMSO control in RD **(E)** and RH30 **(F)** and treatment with 5 µM ATRA, GSK343 or in combination relative to DMSO control in RD **(G)** and RH30 **(H)**. Data shown are mean values from at least 2 independent experiments; bars, SD. One Way ANOVA was used to analyse statistical significance compared to single agent (* p < 0.05, ** p < 0.01, *** p < 0.001, **** p < 0.001.

**Figure S7.**
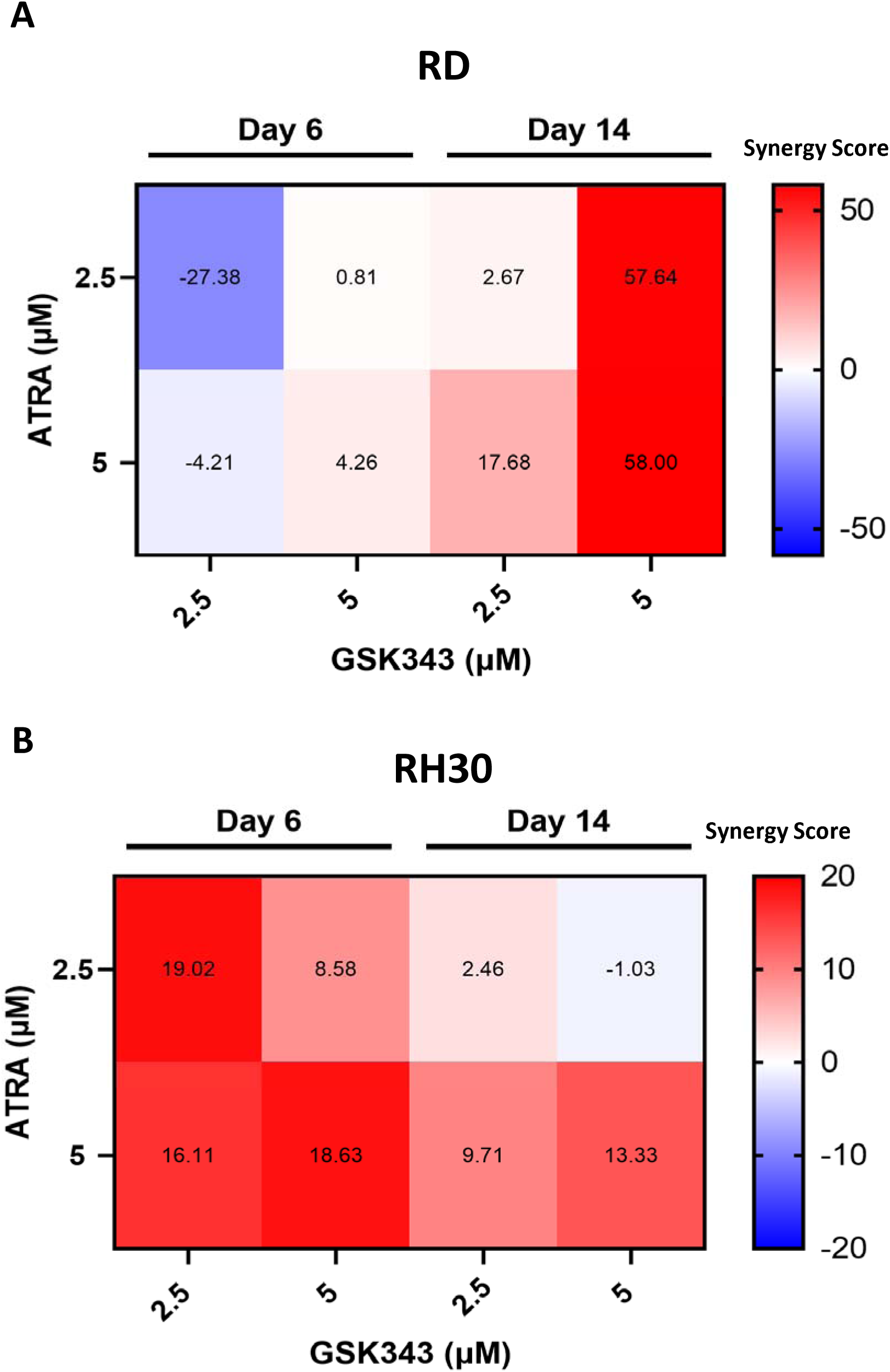
Excess over Bliss calculations of 3D RD (A) and RH30 MCTS (B) treated with combinations of ATRA and GSK343. The predicted Bliss response was subtracted from the experimentally observed inhibition. Scores that are > 0 represent synergy and < 0 represents antagonism.

**Figure S8.**
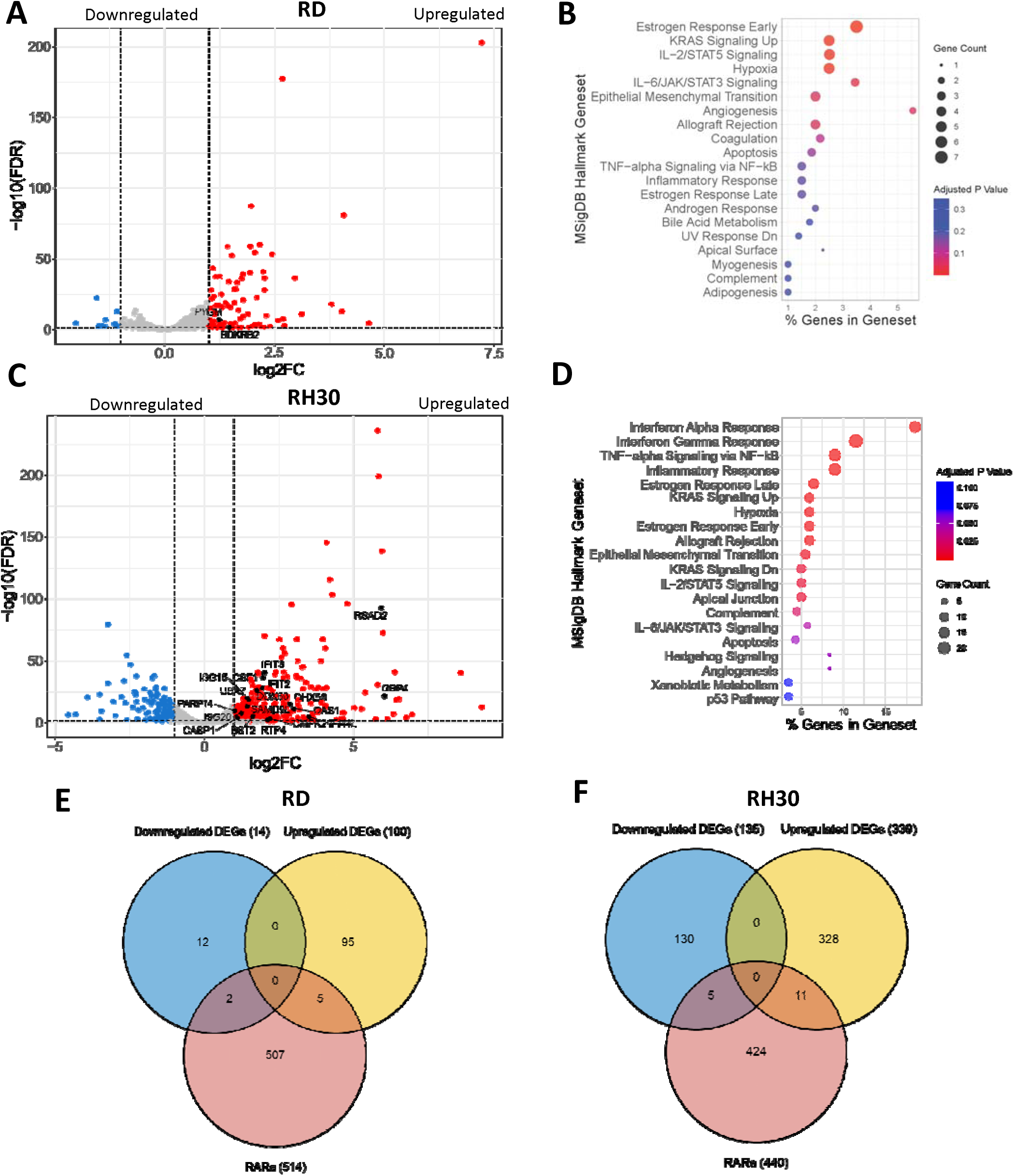
**Integration of RNA-seq and ChIP-seq data in RMS cells treated with 2.5 µM ATRA compared to DMSO control. (A)** Volcano plot of differentially expressed genes after treatment of RD cells, with upregulated myogenesis-related genes highlighted in black. **(B)** Top 20 most significant MSigDB canonical pathways using the upregulated genes from the DEG analysis in RD cells treated with 2.5 µM ATRA. **(C)** Volcano plot of differentially expressed genes after treatment of RH30 cells, with upregulated genes in the canonical interferon α response pathway highlighted in black. **(D)** Top 20 most significant MSigDB canonical pathways using the upregulated genes from the DEG analysis in RH30 cells treated with 2.5 µM ATRA. Venn diagram showing the overlap of number of DEGs with RARα peaks identified from ChIP-seq in **(E)** RD and **(F)** RH30 cells. For all volcano plots, red dots represent

**Figure S9.**
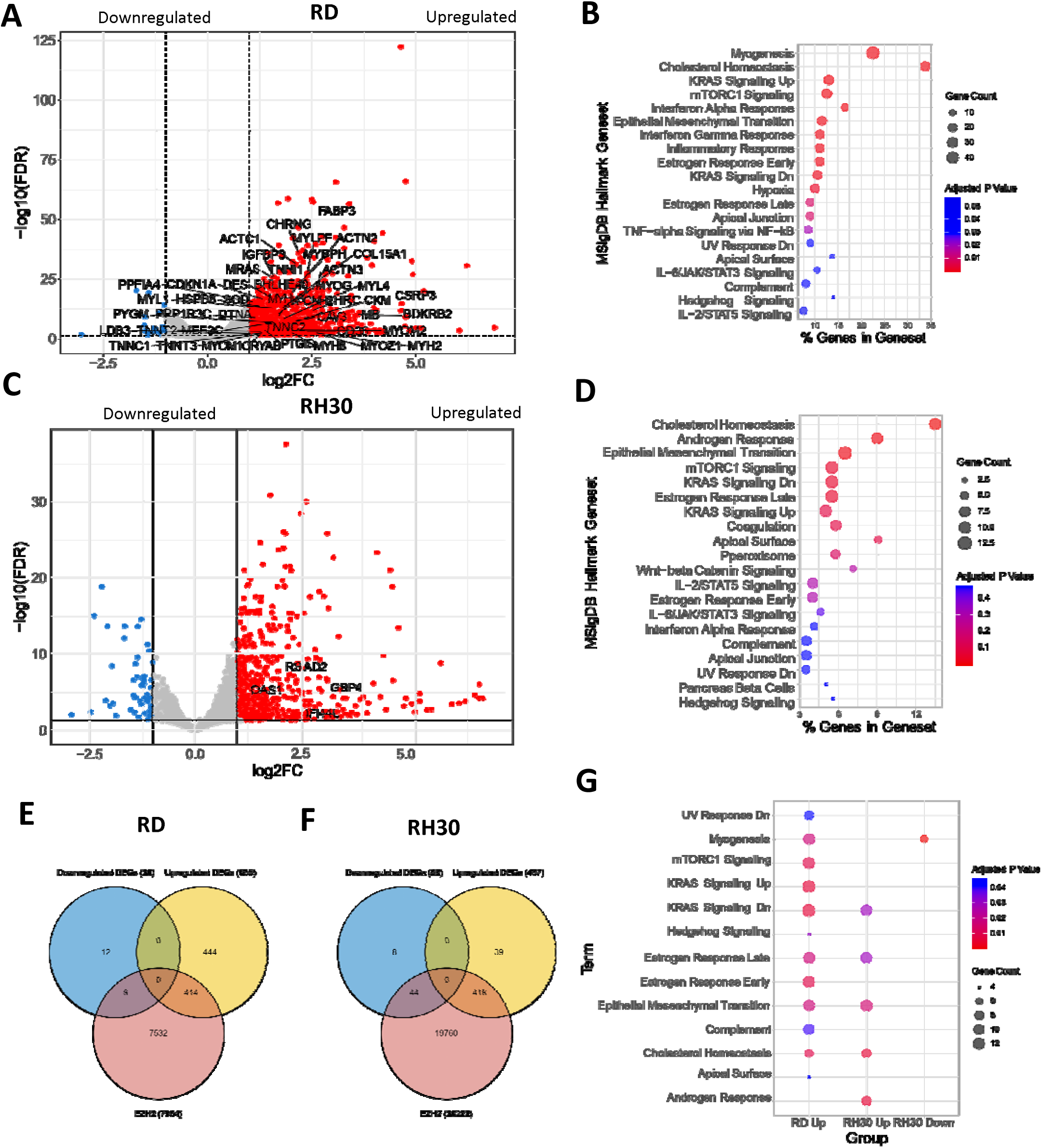
**Figure 6. Integration of RNA-seq and ChIP-seq data in RMS cells treated with 5 µM GSK343 compared to DMSO control. (A)** Volcano plot of differentially expressed genes after treatment of RD cells, with upregulated myogenesis-related genes highlighted in black **(B)** Top 20 most significant MSigDB canonical pathways using the upregulated genes from the DEG analysis in RD cells treated with 5 µM GSK343. **(C)** Volcano plot of differentially expressed genes after treatment of RH30 cells, with upregulated genes in the canonical interferon α response pathway highlighted in black Top 20 most significant MSigDB canonical pathways using the upregulated genes from the DEG analysis with in RH30 cells treated with 5 µM GSK343. Venn diagram showing the overlap of number of differentially expressed genes (DEGs) with EZH2 peaks identified from ChIP-seq in **(E)** RD and **(F)** RH30 cells. **(G)** Gene ontology analysis of DEGs that overlap with EZH2 peaks. For all volcano plots, red dots represent

**Figure S10.**
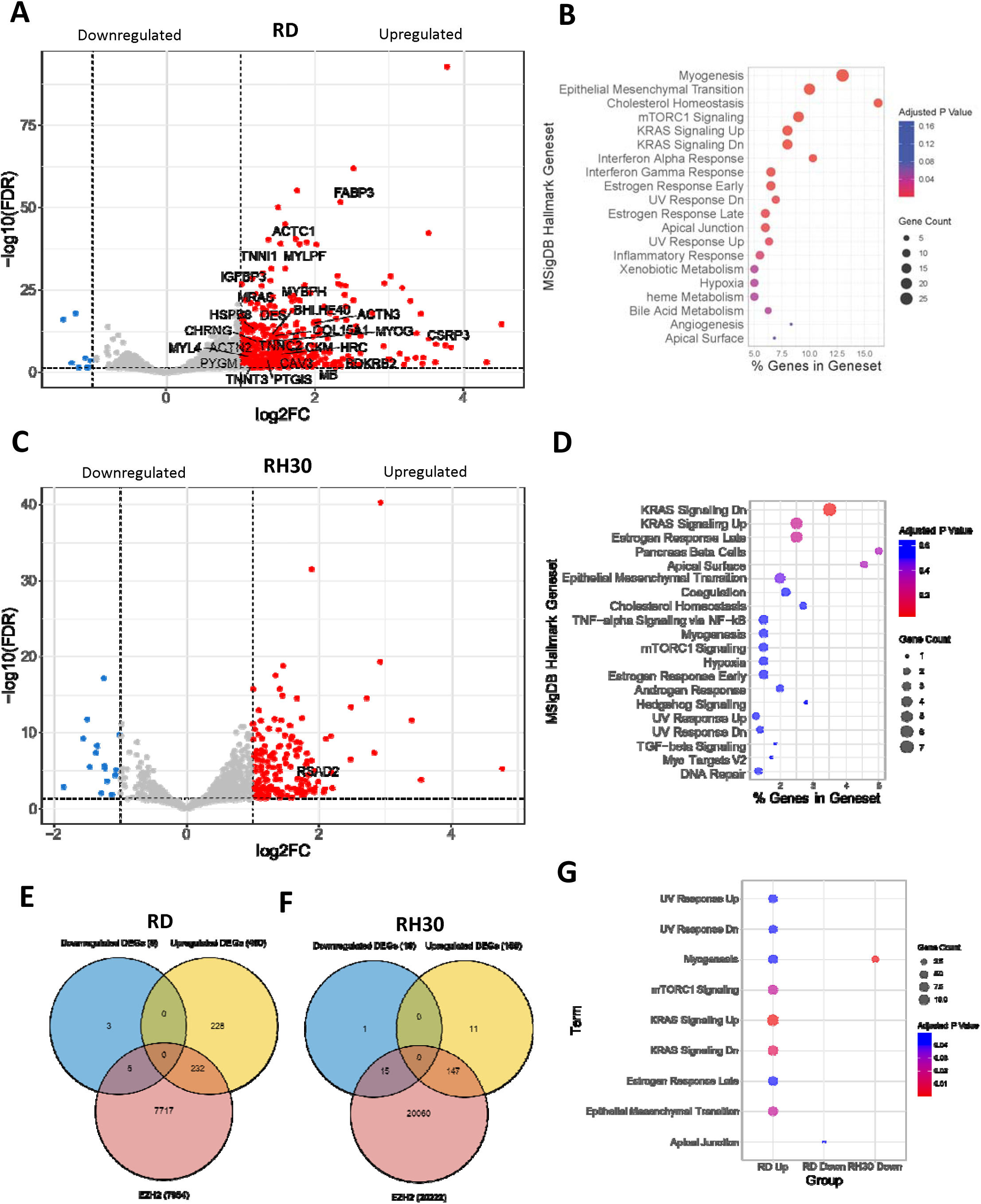
**Integration of RNA-seq and ChIP-seq data in RMS cells treated with 2.5 µM GSK343 compared to DMSO control. (A)** Volcano plot of genes involved in myogenesis in RD cells after treatment. **(B)** Gene ontology analysis of processes that are differentially regulated in RD cells treated with 2.5 µM GSK343. **(C)** Volcano plot of genes involved in the interferon α response pathway in RH30 cells after treatment. **(D)** Gene ontology analysis of processes that are differentially regulated in RH30 cells treated with 2.5 µM GSK343. Venn diagram showing the overlap of number of differentially expressed genes (DEGs) with EZH2 peaks identified from ChIP-seq in **(E)** RD and **(F)** RH30 cells. **(G)** Gene ontology analysis of DEGs that overlap with EZH2 peaks.

**Figure S11.**
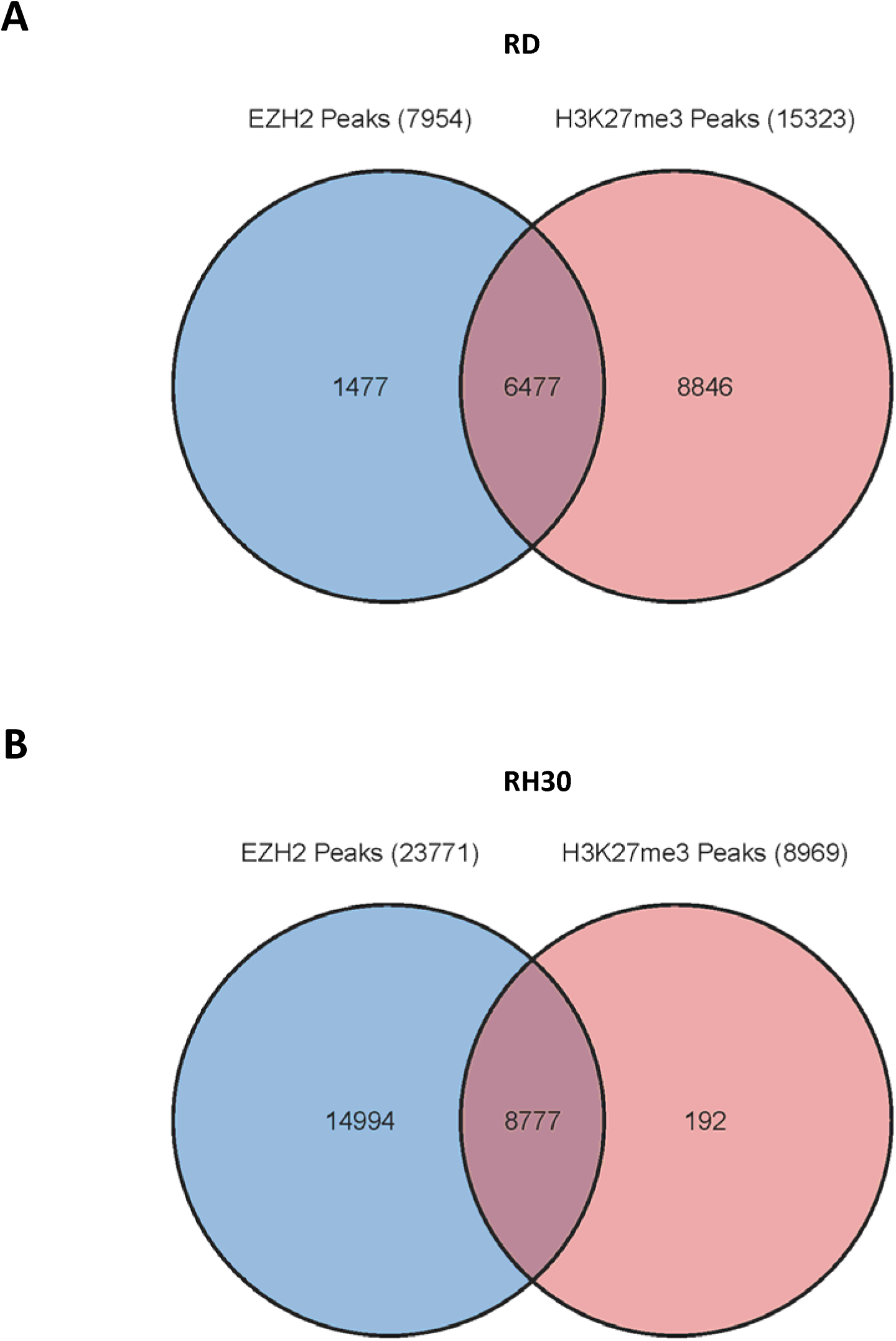
Venn diagram showing the overlap of EZH2 and H3K27me3 peaks from ChIP-seq in (A) RD and (B) RH30 cells.

**Figure S12.**
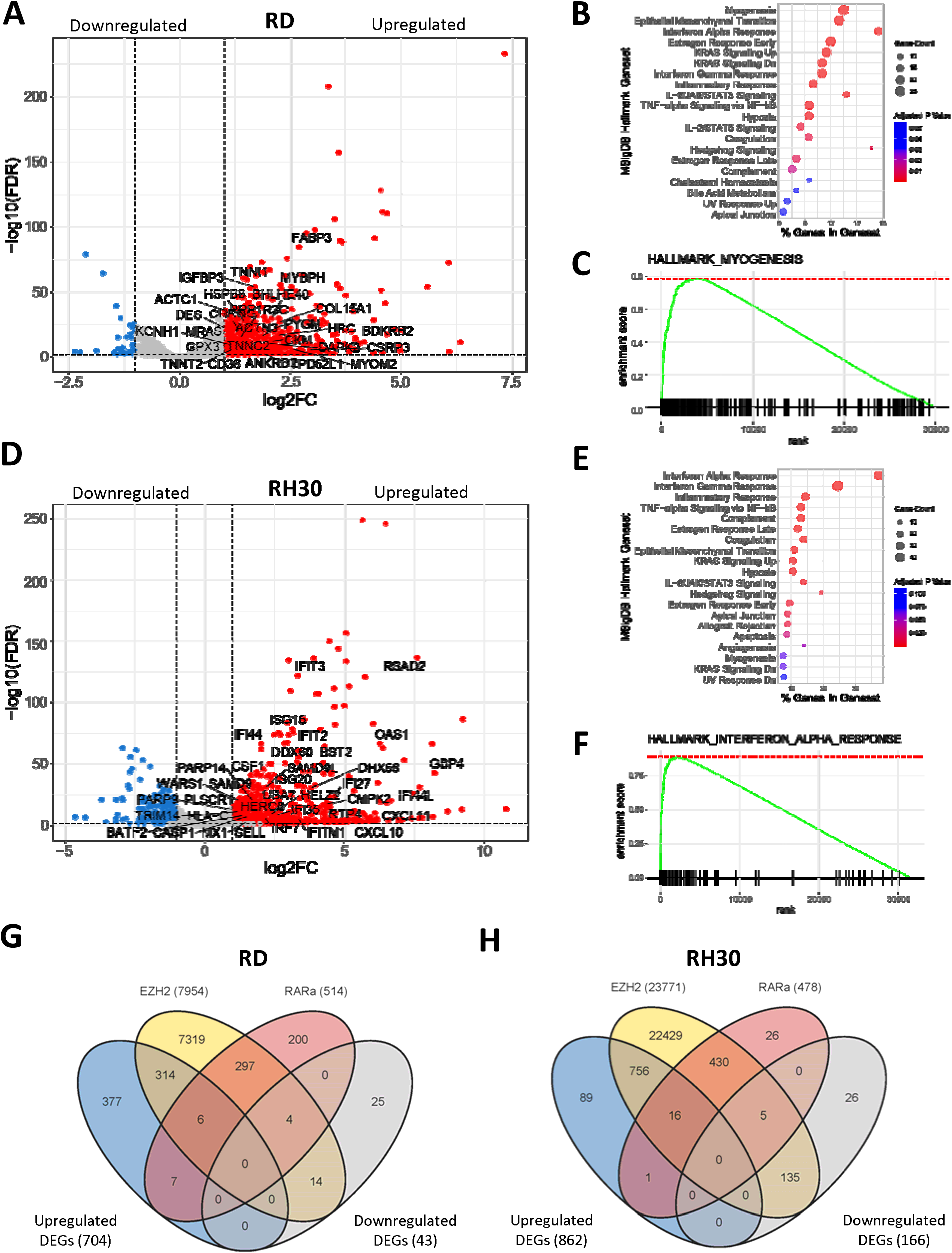
**Integration of RNA-seq and ChIP-seq data in RMS cells treated with a combination of 2.5 µM ATRA and 2.5 µM GSK343 compared to DMSO control (A)** Volcano plot of genes involved in myogenesis in RD cells after treatment. **(B)** Gene ontology analysis of processes that are differentially regulated in RD cells treated with combination of 2.5 µM ATRA and 2.5 µM GSK343. **(C)** GSEA show positive enrichment of myogenesis genes in RD after combination treatment. **(D)** Volcano plot of genes involved in the interferon α response pathway in RH30 cells after treatment. **(E)** Gene ontology analysis of processes that are differentially regulated in RH30 cells treated with combination of 2.5 µM ATRA and 2.5 µM GSK343. **(F)** GSEA show positive enrichment of genes involved in the interferon α response in RH30 after combination treatment. Venn diagram showing the overlap of number of differentially expressed genes (DEGs) with EZH2 and RARα peaks identified from ChIP-seq in **(G)** RD and **(H)** RH30 cells.

**Figure S13.**
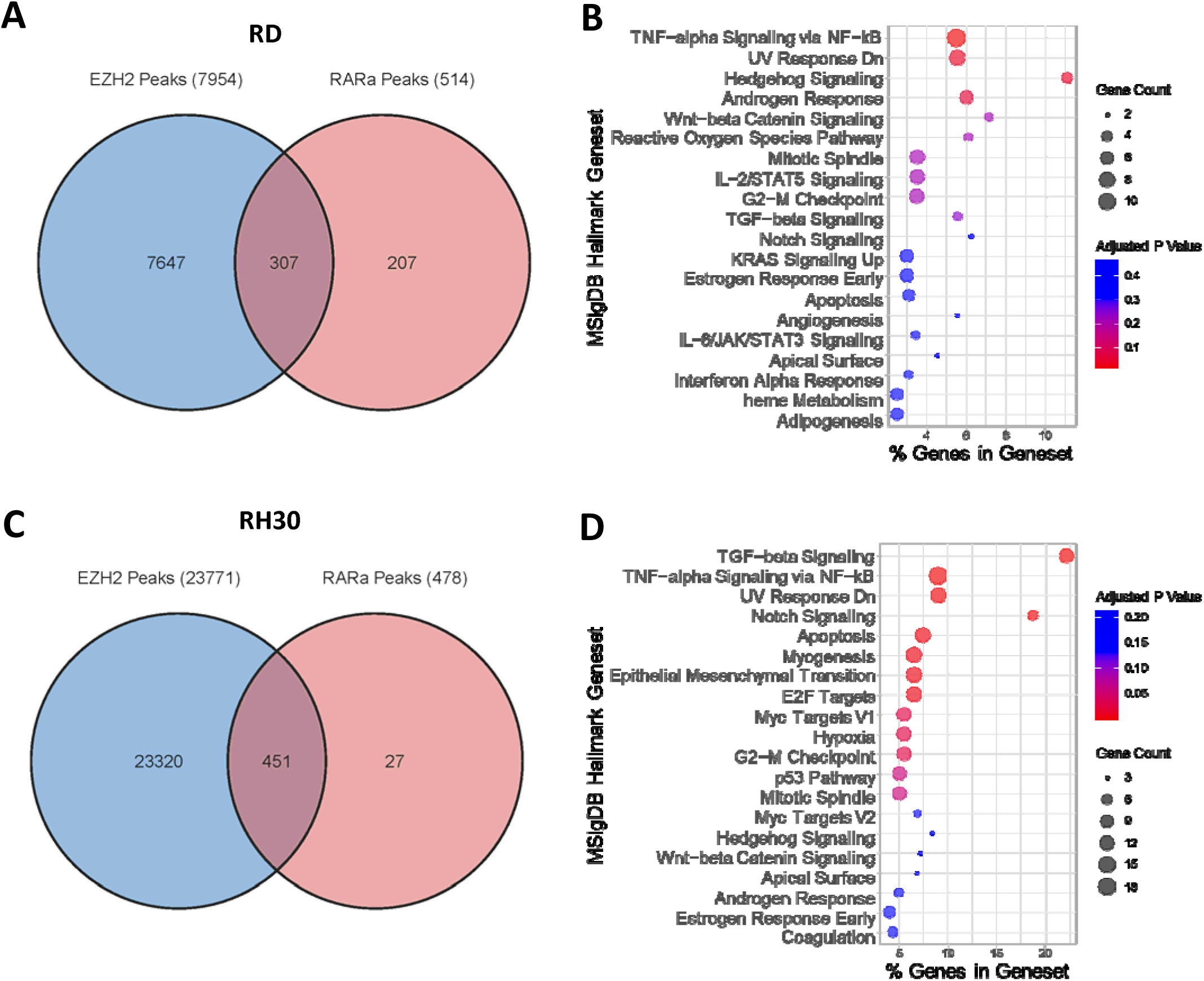
Venn diagram showing the overlap of EZH2 and RARα peaks from ChIP-seq in **(A)** RD cells and **(B)** gene ontology pathway analysis. Venn diagram showing the overlap of EZH2 and RARα peaks from ChIP-seq in **(C)** RH30 cells and **(D)** gene ontology pathway analysis.

**Figure S14.**
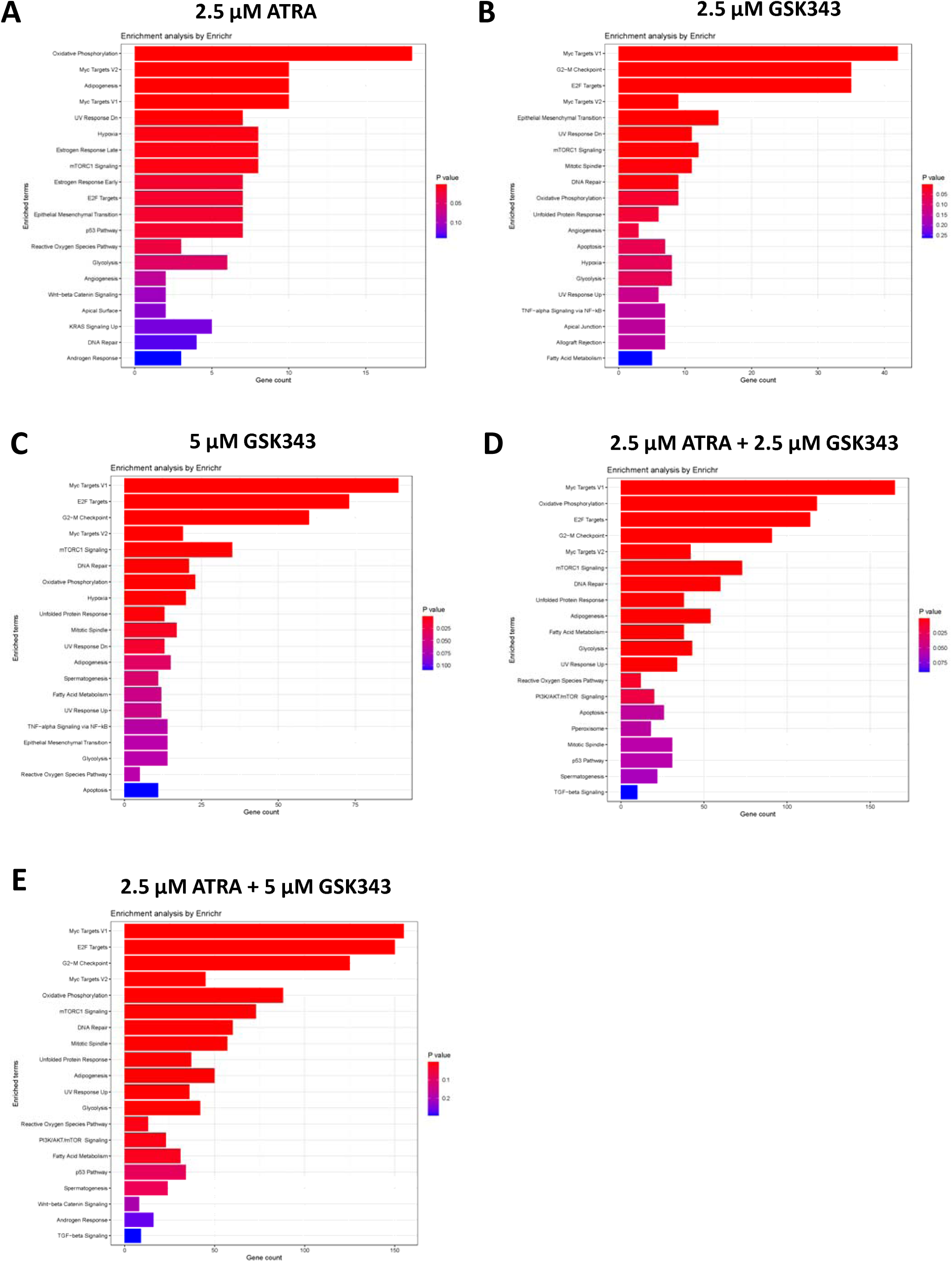
Top 20 most significant MSigDB canonical pathways using the downregulated genes from the DEG analysis in RD cells treated with **(A)** 2.5 µM ATRA **(B)** 2.5 µM GSK343 **(C)** 5 µM GSK343 **(D)** 2.5 µM ATRA + 2.5 µM GSK343 and **(E)** 2.5 µM ATRA + 5 µM GSK343.

**Figure S15.**
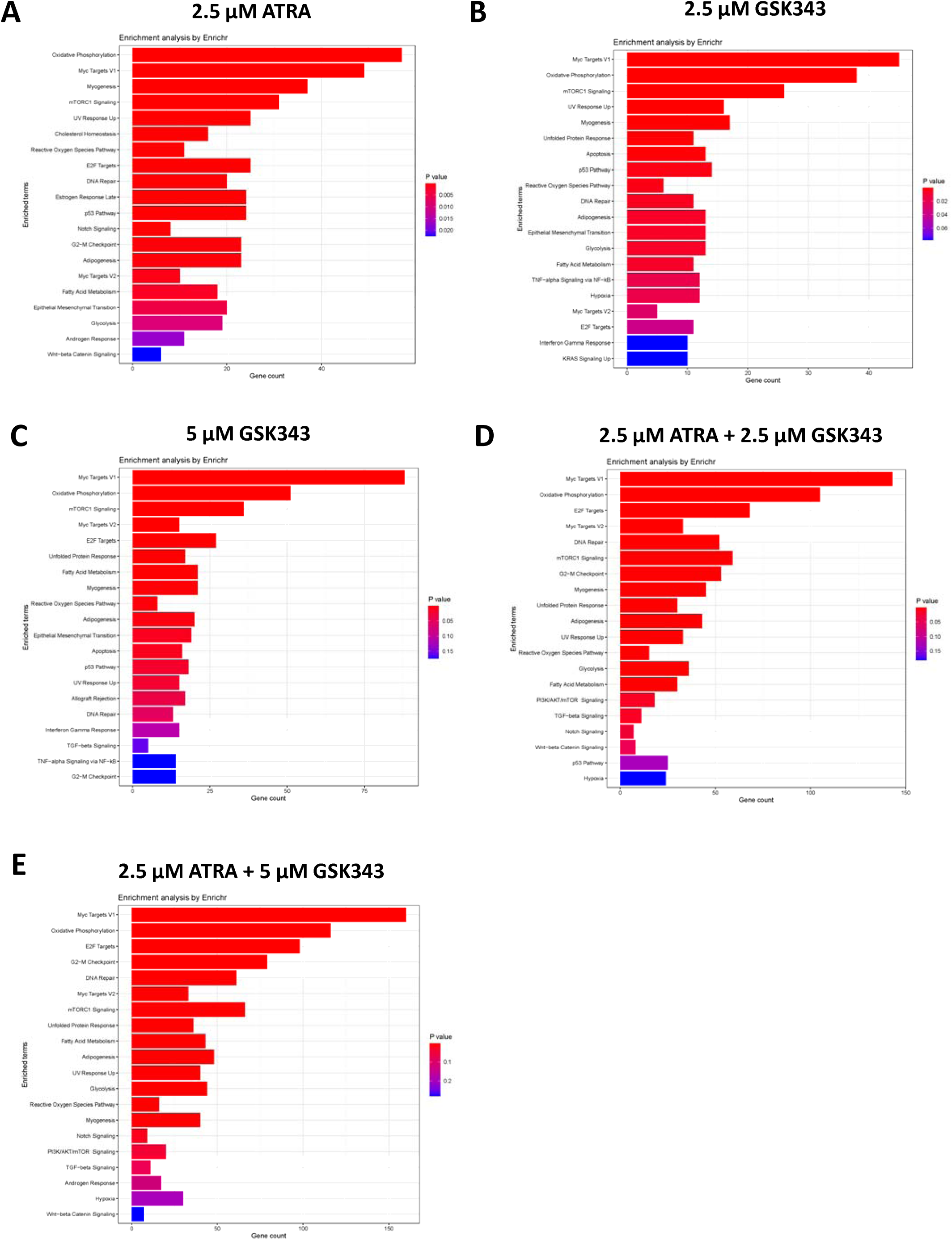
Top 20 most significant MSigDB canonical pathways using the downregulated genes from the DEG analysis in RH30 cells treated with **(A)** 2.5 µM ATRA **(B)** 2.5 µM GSK343 **(C)** 5 µM GSK343 **(D)** 2.5 µM ATRA + 2.5 µM GSK343 and **(E)** 2.5 µM ATRA + 5 µM GSK343.

**Figure S16.**
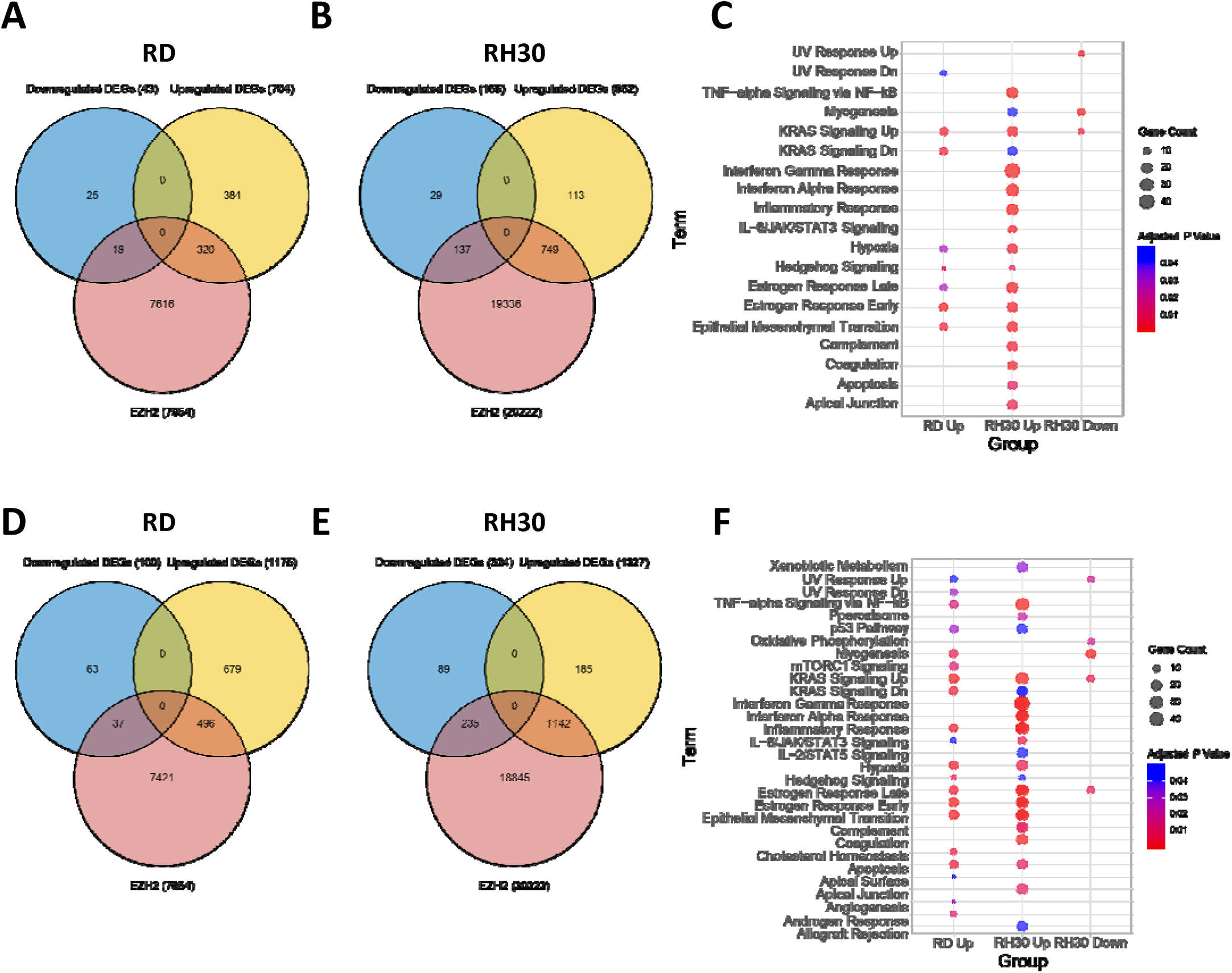
Venn diagram showing the overlap of number of differentially expressed genes (DEGs) with EZH2 peaks identified from ChIP-seq in **(A)** RD and **(B)** RH30 cells treated with combination of 2.5 µM GSK343 and 2.5 µM ATRA. **(C)** Gene ontology analysis of DEGs that overlap with EZH2 peaks after 2.5 µM GSK343 and 2.5 µM ATRA combination treatment. Venn diagram showing the overlap of number of DEGs with EZH2 peaks identified from ChIP-seq in **(D)** RD and **(E)** RH30 cells treated with combination of 5 µM GSK343 and 2.5 µM ATRA. **(F)** Gene ontology analysis of DEGs that overlap with EZH2 peaks after 2.5 µM GSK343 and 2.5 µM ATRA combination treatment.

## Notes

### Competing Interest Statement

The authors have declared no competing interest.

